# Optogenetic delivery of trophic signals in a genetic model of Parkinson’s disease

**DOI:** 10.1101/2020.08.06.238816

**Authors:** Álvaro Inglés-Prieto, Nikolas Furthmann, Samuel Crossman, Nina Hoyer, Meike Petersen, Vanessa Zheden, Julia Biebl, Eva Reichhart, Attila György, Daria Siekhaus, Peter Soba, Konstanze F. Winklhofer, Harald Janovjak

## Abstract

Optogenetics has been harnessed to shed new mechanistic light on current therapies and to develop future treatment strategies. This has been to date achieved by the correction of electrical signals in neuronal cells and neural circuits that are affected by disease. In contrast, the optogenetic delivery of trophic biochemical signals, which support cell survival and thereby may modify progression of degenerative disorders, has never been demonstrated in an animal disease model. Here, we reengineered the human and *Drosophila melanogaster* REarranged during Transfection (hRET and dRET) receptors to be activated by light, creating one-component optogenetic tools termed Opto-hRET and Opto-dRET. Upon blue light stimulation, these receptors robustly induced the MAPK/ERK proliferative signaling pathway in cultured cells. In PINK1^B9^ flies that exhibit loss of PTEN-induced putative kinase 1 (PINK1), a kinase associated with familial Parkinson’s disease (PD), light activation of Opto-dRET suppressed mitochondrial defects, tissue degeneration and behavioral deficits. In human cells with PINK1 loss-of-function, mitochondrial fragmentation was rescued using Opto-dRET *via* the PI3K/NF-кB pathway. Our results demonstrate that a light-activated receptor can ameliorate disease hallmarks in a genetic model of PD. The optogenetic delivery of trophic signals is cell type-specific and reversible and thus has the potential to overcome limitations of current strategies towards a spatio-temporal regulation of tissue repair.

**Significance Statement:** The death of physiologically important cell populations underlies of a wide range of degenerative disorders, including Parkinson’s disease (PD). Two major strategies to counter cell degeneration, soluble growth factor injection and growth factor gene therapy, can lead to the undesired activation of bystander cells and non-natural permanent signaling responses. Here, we employed optogenetics to deliver cell type-specific pro-survival signals in a genetic model of PD. In *Drosophila* and human cells exhibiting loss of the PINK1 kinase, akin to autosomal recessive PD, we efficiently suppressed disease phenotypes using a light-activated tyrosine kinase receptor. This work demonstrates a spatio-temporally precise strategy to interfere with degeneration and may open new avenues towards tissue repair in disease models.

## Introduction

Biology occurs over a wide range of time and length scales, from milliseconds and nanometers for protein folding, to days and centimeters for organism development. In recent years, powerful research methods have been developed that permit the manipulation of biological processes on even the smallest length and shortest time scales. In optogenetics, natural or reengineered photoreceptors are expressed in genetically defined cell populations to optically activate or inhibit, e.g., neuronal action potential firing or cell signaling. The use of light provides unprecedented precision in space and time as a way to answer previously unresolvable questions in a multitude of disciplines, including microbiology, cell/developmental biology, synthetic biology, and neuroscience. In particular, spatio-temporally precise perturbation of selected cells in intact organisms can reveal cause-consequence-relationships that are a critical determination for understanding central nervous system function or animal development (1–3). Optogenetics also provides access to the reversible and rapid activation of cell signaling pathways that is required for dissection of their dynamic properties (4, 5) and for development of new drug discovery platforms (6). Inspired by these successes, optogenetics is continuously translated into new research areas, including disease mechanism and therapy.

Shortly after its inception, optogenetics was beginning to be employed in the study of neural circuits that are known to be affected by neurological and neurodegenerative disorders, including spinal cord injury, stroke and Parkinson’s disease (PD) (7, 8). In this field, optogenetics has shed new light on the mechanisms of currently utilized therapies (e.g., deep brain stimulation in PD) or therapies of the future (e.g., stem cell-based tissue regeneration) (9, 10). This work was followed by the development of light-gated prosthetic approaches in which a genetically introduced photoreceptor senses either natural light, e.g. for vision restoration (11), or light from a prosthetic source, e.g. for heart or skeletal muscle pacing (12, 13). Notably, these pioneering studies harnessed optogenetics to excite or inhibit electrical signals through regulated ion flow ions across the cell membrane. In apparent contrast, the optogenetic delivery of trophic signals, which support cell survival and are central to treatment strategies in a variety of degenerative disorders, has never been demonstrated in a disease model. It is unclear if this is feasible as hypo- or hyperactivity of pro-survival pathways is linked to undesired cellular outcomes (see below).

We and others have recently engineered light-activated variants of key signaling proteins that now provide a basis for the optogenetic delivery of trophic signals. Particular success was reported for receptor tyrosine kinases (RTKs) (14–18). RTKs are expressed in virtually all human cell types and respond to growth factors (GFs) with conformational changes and/or oligomerization state changes that result in receptor *trans*-phosphorylation. *Trans*-phosphorylation is then followed by recruitment of intracellular secondary messengers in, e.g., the mitogen-activated protein kinase/extracellular signal-regulated kinase (MAPK/ERK) or phosphatidylinositol-3 kinase/AKT (PI3K/AKT) signaling pathways. Because of their ability to activate these proliferative and pro-survival pathways, RTKs are prime targets in several neurodegenerative disorders. In the context of PD, the RET RTK (19) has been intensively investigated in both preclinical and clinical studies. hRET is activated by glial cell line-derived neurotrophic factor (GDNF) family ligands (GFLs; these are GDNF, neurturin, artemin, and persephin) that bind GDNF family receptor α (GFRα) co-receptors (GFR α1-4) to recruit dimeric RET into a ternary complex (**Figure 1A**). GFLs are linked to the development and maintenance of dopaminergic (DA) midbrain neurons and have been pursued as disease-modifying agents in PD, either by local injection or by gene delivery using adeno-associated viruses (20, 21). Despite initial success in animal models, outcomes in clinical trials were limited (22, 23), which was attributed to difficulties in GFL delivery, limited responsiveness of the targeted DA neurons and advanced PD in some of the recruited patients. In addition, there are concerns that the continuous delivery of GFLs can lead to counter-productive compensatory effects (24–27). These observations and considerations have highlighted a need for methods that can control the GFL-RET-axis in a reversible and more precise manner (28, 29).

**Figure 1.**
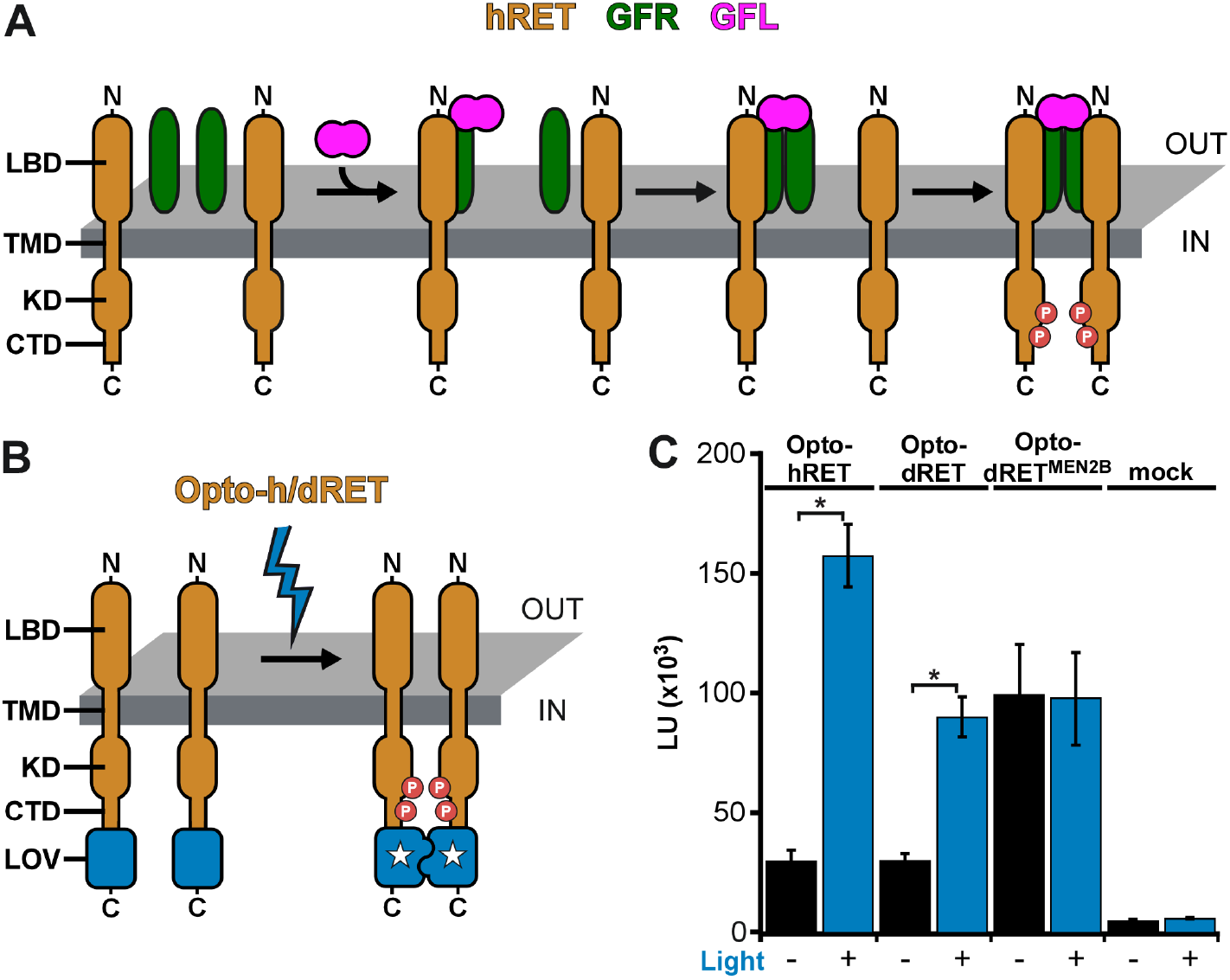
Engineering of light-activated RET receptors. (**A**) hRET and dRET consist of an extracellular ligand-binding domain (LBD), single-span transmembrane domain (TMD) and intracellular domain (KD: kinase domain, CTD: C-terminal tail domain). Activation by GFL and GFRα was shown to result in the formation of a human ternary complex (binding model of Schlee *et al.* (39)). (**B**) In light-activated Opto-h/dRET, the LOV domain of the AUREOCHROME1 photoreceptor of *V. frigida* is incorporated at the receptor C-terminus. (**C**) MAPK/ERK pathway activation in response to blue light (I = 250 μW/cm^2^, λ = 470 nm) for HEK293 cells transfected with *Opto-hRET*, *Opto-dRET* or *Opto-dRET^MEN2B^*. Light units (LU; mean ± SD) for dark treated cells and illuminated cells are given (n = 9 to 18, three independent experiments, t-test, *: p<.0001).

Here, we explored optogenetics as a means for delivery of trophic signals in a genetic model of PD. We first reengineered full-length hRET and its *Drosophila* orthologue dRET (30–32) to be activated by light in optogenetic tools termed Opto-hRET and Opto-dRET. We then showed that temporally precise dRET activation *in vivo* can be used to induce degeneration in a tissue sensitive to ectopic RTK signaling. Optogenetic delivery of RET signals was then successfully applied in a genetic fly model of PD. Mutations in the *PINK1* gene are linked to autosomal recessive PD (33–35), and *Drosophila* has been shown to be a suitable model to study consequences of PINK1 loss-of-function (36–38). We suppressed *Drosophila* phenotypes associated with PINK1 deficiency and identified the involved downstream signaling pathways in a human cellular model. This work demonstrates the use of optogenetics as a cell-type specific and remote controlled method to exert beneficial trophic effects of in a genetic disease model.

## Results

### Light-activated hRET and dRET receptors

hRET assembles in dimers in the activated ternary GFL_2_-GFRα_2_-RET_2_ complex (39, 40) (**Figure 1A**) and forced dimerization by mutations or synthetic binding domains has been shown to induce signaling of hRET (41, 42) and dRET (43, 44). Based on these observations, we converted hRET and dRET into optogenetic tools by incorporating a light-activated dimerization switch. To achieve this switch, we utilized the light-oxygen-voltage-sensing (LOV) domain of the AUREOCHROME1 photoreceptor from the yellow-green algae *Vaucheria frigida* (AU1-LOV) (45) (**Figure 1B**). AU1-LOV is a member of the large LOV domain superfamily and responds to blue light with formation of a symmetric homodimer (46) (**Figure S1**). AU1-LOV is smaller than other photoreceptors commonly used in optogenetics (145 aa in length; this corresponds to ~a third of the length of cryptochromes or phytochromes) (47, 48) and relaxes slower than many other LOV domains from the light-activated ‘lit’ state (that is predominantly dimeric) to the dark-adapted state (that is predominantly monomeric; relaxation time constant ~600 s) (49). We and others have shown that small size and slow cycling make AU1-LOV well suited for assembly and activation of membrane receptors (14, 16, 17, 50). We placed AU1-LOV at the far C-terminus of the RET receptors because fluorescent proteins (FPs) were previously incorporated at this site without negative impact on receptor signaling or trafficking (51, 52). To functionally test the generated Opto-hRET and Opto-dRET, we took advantage of the fact that *Drosophila* RTKs can couple to the mammalian MAPK/ERK pathway *via* Ras (44). We and others utilize human embryonic kidney 293 (HEK293) cells for testing new optogenetic methods because these cells do not exhibit native light-induced signaling events. Using transcriptional reporters (14, 47), we found robust induction of MAPK/ERK signaling upon blue light stimulation of HEK293 cells transfected with Opto-hRET and Opto-dRET (intensity (I) = 250 μW/cm^2^, wavelength (λ) = 470 nm) (**Figure 1C**). Whilst Opto-hRET activated transcriptional responses more strongly than Opto-dRET, Opto-dRET activation levels were comparable to those reached by the Opto-dRET^MEN2B^ variant (**Figure 1C**). Opto-dRET^MEN2B^ contains a Met to Thr gain-of-function substitution in the kinase domain that was discovered in multiple endocrine neoplasia (MEN) Type 2B as causative for potent receptor hyperactivation in the absence of GFLs (43, 53). These results show that signaling activity can be induced by blue light through Opto-h/dRET receptors.

### Opto-dRET function in vivo

We next tested if Opto-dRET can be applied *in vivo* to conduct a temporally-targeted gain-of-function experiment (**Figure 2A**). We choose the *Drosophila* retina for this experiment because RTKs and their downstream pathways are tightly regulated during its morphogenesis, and because RTK hyperactivation during retina development results in marked phenotypes. For instance, two RTKs, the *Drosophila* epidermal growth factor receptor (DER) and Sevenless, orchestrate retinal cell growth, differentiation and regulated death (54, 55). These RTKs act in late larval and early pupal stages to form the fourteen cells that compose each ommatidium unit eye and the ommatidial lattice (56). Hyperactivation of RTK signaling during these stages has been shown to result in irregular ommatidia size and spacing leading to a disrupted tissue pattern termed ‘roughening’ (57). We generated transgenic flies in which the *Opto-dRET* gene is placed downstream of five *UAS* elements (**Figure S2**). We also generate analogous flies expressing the constitutively-active Opto-dRET^MEN2B^. We then targeted Opto-dRET or Opto-dRET^MEN2B^ to the retina using the *GMR-GAL4* driver, which induces expression in cells located posterior of a morphogenetic furrow that sweeps in anterior direction to initiate mitosis and cell differentiation (55). In Opto-dRET^MEN2B^ flies, scanning electron microscopy (SEM) revealed a marked rough retina phenotype (compare **Figure 2B** and **C**). Roughening was previously observed in flies expressing dRET^MEN2B^ (43), and based on the severe outcome observed for Opto-dRET^MEN2B^ we concluded that AU1-LOV attachment does not negatively impact receptor function. We next illuminated control flies and Opto-dRET flies in a time window that captures ommatidia and lattice formation (from third instar larva through to the second day after pupariation; I = 385 μW/cm^2^, λ = 470 nm; **Figure 2A**). In control flies without Opto-dRET, we did not observe light-induced roughening indicating that light alone did not have an effect on the retina (compare **Figure 2D** and **E**; in agreement with previous studies, we observed mild phenotypes upon GAL4 expression with the *GMR* driver (58)). In apparent contrast, we found that light stimulation resulted in a marked increase in roughening in Opto-dRET flies (compare **Figure 2F** and **G**). To quantify this effect, we manually metered in each retina image the ‘fused area’ (the area without identifiable ommatidia) and also applied computational methods to count individual ommatidia (~600 ommatidia can be assigned in our frontal view images) as two measures of tissue integrity. We found that upon illumination the fused area increased and the number of identified structures decreased specifically in the illuminated Opto-dRET flies (**Figure 2H** and **I**). For these and control flies, we also determined lattice regularity, which is defined as the ratio of the mean and the standard deviation (SD) of the ommatidia nearest-neighbor distance (NND) distribution (59). Regularity decreased from 3.98 ± 0.39 in WT flies to 2.15 ± 0.55 in illuminated Opto-dRET flies, and these values are indicative of near-perfect regularity and near-random assembly, respectively (60). The potent effects induced by Opto-dRET upon light stimulation and the lack of light responses in the absence of Opto-dRET establish the suitability of this optogenetic approach to modify tissue behavior *in vivo*.

**Figure 2.**
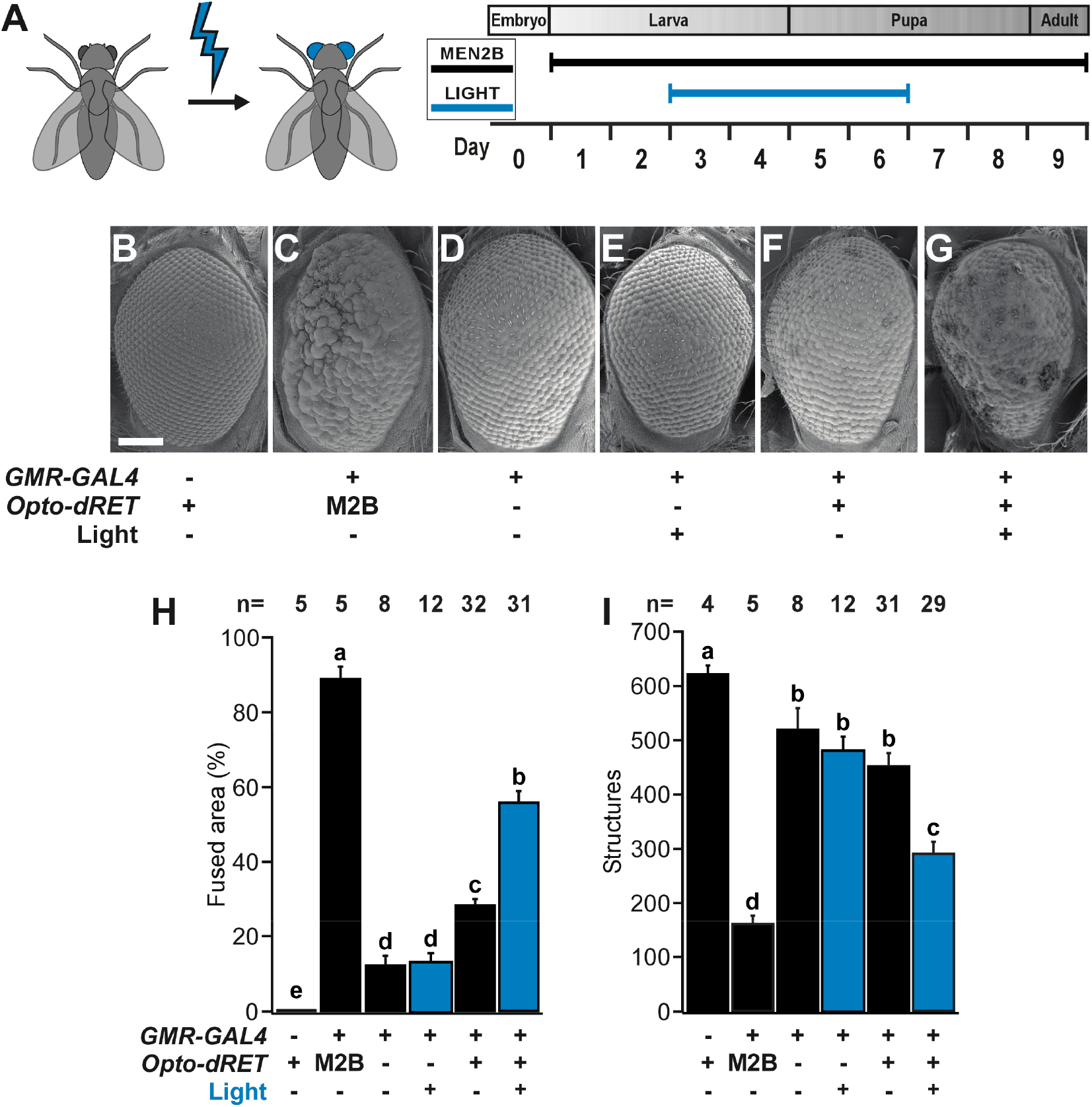
Induction of retina roughening and phenotype quantification. (**A**) Developmental time window targeted by light in retina experiments. (**B-G**) Representative retina SEM images. Scale bar: 0.1 mm. (**H** and **I**) Quantification of rough retina phenotypes of one-day old flies as fused area and the number of structures identified. “M2B” denotes Opto-dRET^MEN2B^. For H and I, mean ± SD for the indicated number of flies is given (at least three independent experiments). Means sharing the same label are not significantly different (ANOVA/Bonferroni corrected t-tests, p>.04). Light intensity was 385 μW/cm^2^.

### Suppression of defects in a genetic model of PD

With Opto-dRET in hand, we went on to explore if defects in a genetic disease model can be ameliorated using optogenetics (**Figure 3A**). PINK1 is a Ser/Thr kinase that localizes to mitochondria and supports their integrity and function. Loss-of-function mutations and dominant negative mutations in the *PINK1* gene are associated with autosomal recessive PD (33–35). In *Drosophila*, loss of X-linked *PINK1* leads to a striking phenotype, including tissue degeneration, locomotor defects and disruption of mitochondrial structure and function (61–64). To test if optogenetics can suppress phenotypes associated with PINK1 deficiency, we expressed Opto-dRET in indirect flight muscles (IFMs) of PINK1^B9^ flies using the *MEF2-GAL4* driver (65). IFMs are frequently studied in *Drosophila* models of PD and PINK1 loss-of-function leads to a marked ‘crushed’ thorax phenotype and reduced locomotion. We first compared PINK1^B9^ flies to Opto-dRET PINK1^B9^ flies that were not illuminated. Similar penetrance of thoracic defects (58 and 61% of flies exhibited a crushed thorax, respectively) shows that the engineered optogenetic receptor did not affect the phenotype in the absence of light (**Figure 3B**). When proceeding to light stimulation, we took into consideration that the opaque case and cuticle of pupa and adults may reduce blue light penetration to IFMs. To address this, we first confirmed that AU1-LOV can be activated by light of 1-5 μW/cm^2^ intensity, which corresponds to the product of minimal blue light transmission through the case or cuticle (~0.5% (66, 67)) and the light intensity applied in our light chambers (I = 320 μW/cm^2^; **Figure S3A**). We observed that light of this intensity is indeed sufficient to activate AU1-LOV (**Figure S3B**). We then went on to light stimulate Opto-dRET PINK1^B9^ flies during late pupal stages and adult states (**Figure 3A**) (these stages coincide with the onset of degeneration (64, 68)). Strikingly, we observed phenotype suppression in Opto-dRET PINK1^B9^ flies resulting in only 16% of flies with defects (**Figure 3B**). This result indicates marked improvement in tissue integrity and was comparable to the improvement observed previously upon PINK1 overexpression in the PINK1^B9^ model (Park et al, 2006). We also tested if illumination restored the climbing deficits that accompany PINK1 loss-of-function.

**Figure 3.**
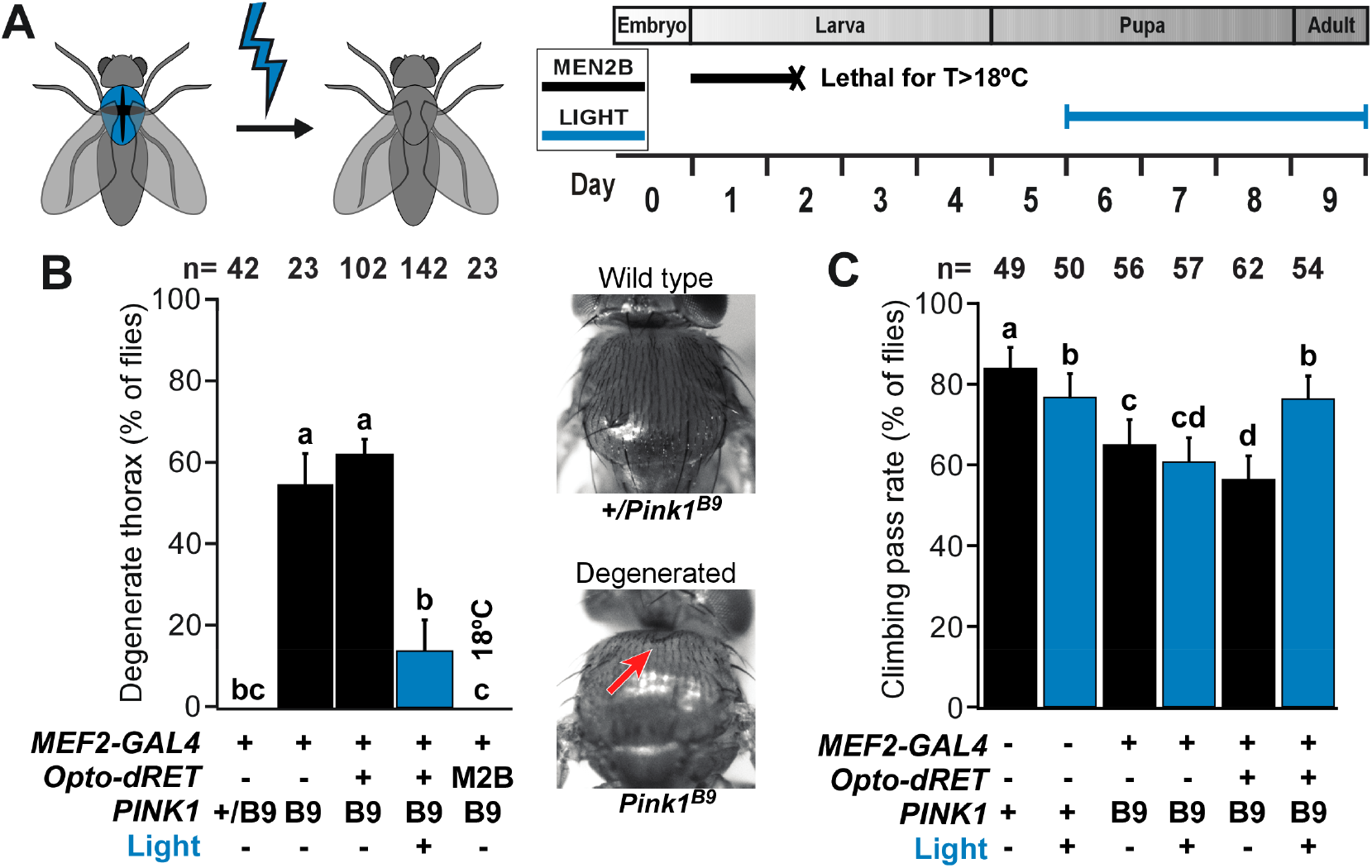
Suppression of thorax defects and locomotion deficits. (**A**) Time window targeted by light in experiments with PINK1^B9^ flies. Illumination of pupal and adult stages prevented lethality observed upon Opto-dRET signaling in earlier stages (e.g., Opto-dRET^MEN2B^ flies were grown at 18°C to prevent lethality during development; see Main Text). Percentage of two-day old flies with a degenerate thorax phenotype. Representative bright field thorax images shown on the right. Hollow thorax is highlighted by the red arrow. (B) Climbing ability of flies. “M2B” denotes Opto-dRET^MEN2B^. *PINK1* “+” denotes the WT gene. For B and C, counts ± SE for the indicated number of flies (n) is given (see Materials and Methods for repetitions in climbing assays). Means sharing the same label are not significantly different (Fisher’s exact test, p>.04). Light intensity was 320 μW/cm^2^.

This was indeed the case with illuminated Opto-dRET flies reaching climbing pass rates similar to those of WT flies (**Figure 3C**).

Mitochondrial dysfunction is a major pathological alteration observed in sporadic and familial PD and also the primary cellular consequence of loss of PINK1. We therefore tested the effect of illumination on mitochondrial function and integrity in Opto-dRET PINK1^B9^ flies. PINK1^B9^ flies exhibited reduced muscle ATP levels and these levels could be restored by Opto-dRET and light stimulation (**Figure 4A**). To examine mitochondrial integrity, we conducted ultrastructure analysis using transmission electron microscopy (TEM). PINK1^B9^ muscles exhibited broadening of the myofibril Z-line and enlarged mitochondria with fragmented cristae (compare **Figure 4B** and **C**). Illumination of Opto-dRET PINK1^B9^ flies was clearly beneficial with a reduced fraction of impaired mitochondria and an increased fraction of mitochondria with WT-like cristae structure (compare **Figure 4D** and **E**) that approached levels of control flies (**Figure 4F**). Overall, these results on the cell- and tissue-level demonstrate optogenetic suppression of consequences of PINK1 loss-of-function in a *Drosophila* model. In these experiments, we took advantage of temporally precise light stimulation to prevent undesired side effects of continuous growth signal delivery, specifically lethality associated with dRET overactivation in muscle at early developmental stages (69).

**Figure 4.**
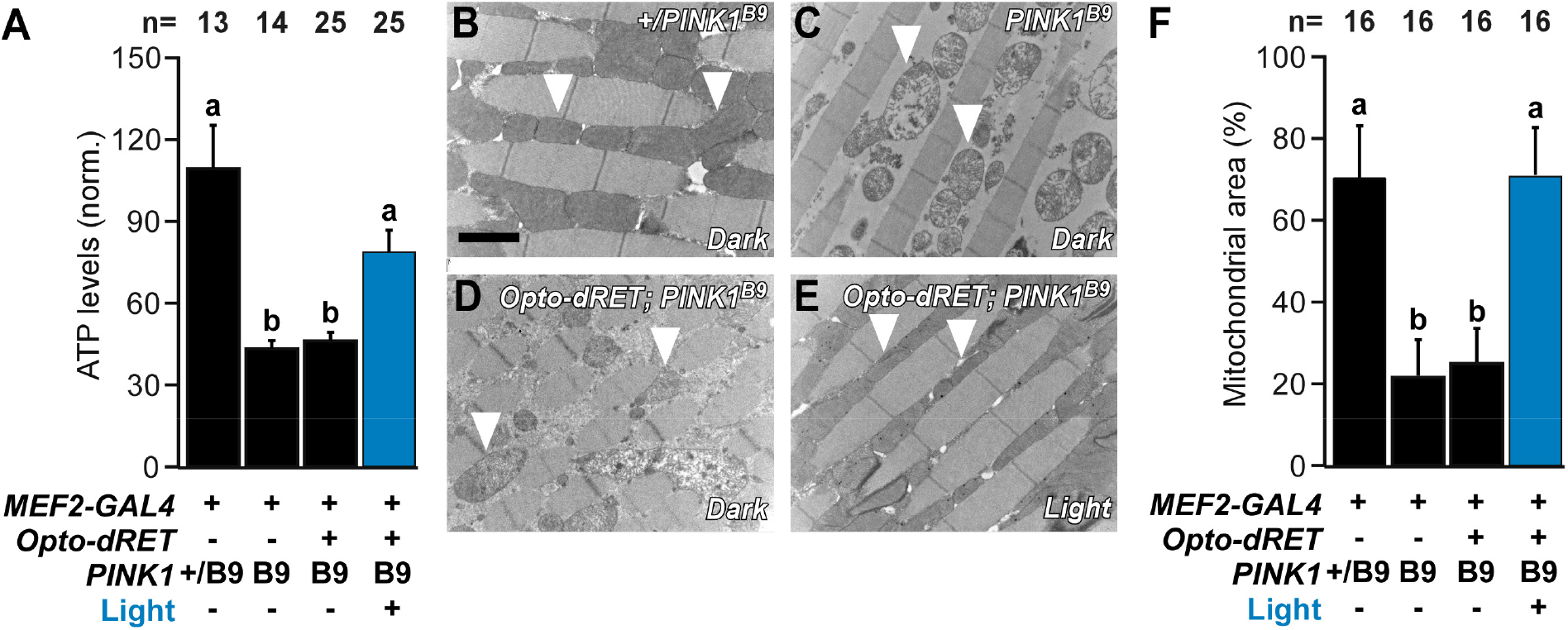
Improved mitochondrial structure and function. (**A**) ATP content in fly thoraces from PINK1^B9^ flies at the indicated conditions. (**B-E**) Representative TEM images of thoracic indirect flight muscles. Arrow heads indicate mitochondria that are either electron dense (B: controls, E: illuminated PINK1^B9^ Opto-dRET flies) or malformed with disintegrated cristae (C: PINK1^B9^ flies, D: PINK1^B9^ Opto-dRET flies in the absence of light). Scale bar: 2 μm. (**F**) Analysis of mitochondrial density in TEM images. “M2B” denotes Opto-dRET^MEN2B^. *PINK1* “+” denotes the WT gene. For A, mean ± SD for the indicated number of flies is given (at least three independent experiments). For F, mean ± SD for the indicated number of micrographs is given (at least three independent experiments). Means sharing the same label are not significantly different (ANOVA/Bonferroni corrected t-tests, p>.04). Light intensity was 320 μW/cm^2^.

### Amelioration of mitochondrial defects in PINK1-deficient human cells

Finally, we tested if light activation of RET signaling can revert defects induced by loss of PINK1 in human cells. We performed these experiments in dopaminergic neuroblastoma SH-SY5Y cells that have been previously applied to study how mutations observed in PD, including those in the *PINK1* gene (70), impact mitochondrial integrity. We transfected the cells with either control siRNA or *PINK1* siRNA as well as expression vectors for Opto-dRET, Opto-dRET^MEN2B^ or an inactive ‘kinase-dead’ (KD) Opto-dRET (Opto-dRET^KD^). Western blot (WB) analysis revealed efficient downregulation of PINK1 levels (**Figure 5A**) and that expression levels of the Opto-dRET variants were comparable (**Figure 5B**). Silencing of the *PINK1* gene resulted in severe mitochondrial defects with ~65% of cells exhibiting fragmented mitochondria (**Figure 5C**, rows 1 and 2, **Figure 5D**, bars 1 and 2). As shown previously, mitochondrial integrity was restored in this model through endogenous RET stimulated with GDNF/GFRα1 for 4 h (69, 71). In this paradigm, the fraction of cells with fragmented mitochondria was reduced to 20%, which is comparable to cells treated with control siRNA (**Figure 5C**, rows 1 and 3, **Figure 5D**, bars 1 and 3). We then analyzed Opto-dRET^MEN2B^ and Opto-dRET cells and found that either expression of Opto-dRET^MEN2B^ or light stimulation of Opto-dRET cells (I = 232 μW/cm^2^, λ = 470 nm, 4 h) rescued mitochondria with similar efficiency (~25% of cells displaying fragmentation; **Figure 5C**, rows 4 to 6, **Figure 5D**, bars 4 to 6). Similarly to the *Drosophila* experiments, no rescue was observed upon Opto-dRET expression in dark conditions, indicating limited basal receptor activity in the absence of the light stimulus (**Figure 5C**, rows 2 and 5, **Figure 5D**, bars 2 and 5). We also verified that the kinase activity of dRET is required for rescue (**Figure 5C**, rows 5 to 8, **Figure 5D**, bars 5 to 8) and that light alone did not influence mitochondrial morphology (**Figure S4**). We also tested which signaling pathways downstream of RET are involved in mediating mitochondrial integrity. Of the main pathways activated by RET, we found that reversion of mitochondrial fragmentation depended on both the PI3K and nuclear factor ‘kappa-light-chain-enhancer’ of activated B-cells (NF-кB) pathway, but not on the MAPK/ERK pathway (**Figure 5E**). This result is in agreement with previous studies showing that the protein network regulated by GFL/RET overlaps with that involved in PINK1 function (69, 71). Collectively, these findings show that beneficial trophic signals can be delivered to a human cellular model of PD using optogenetics.

**Figure 5.**
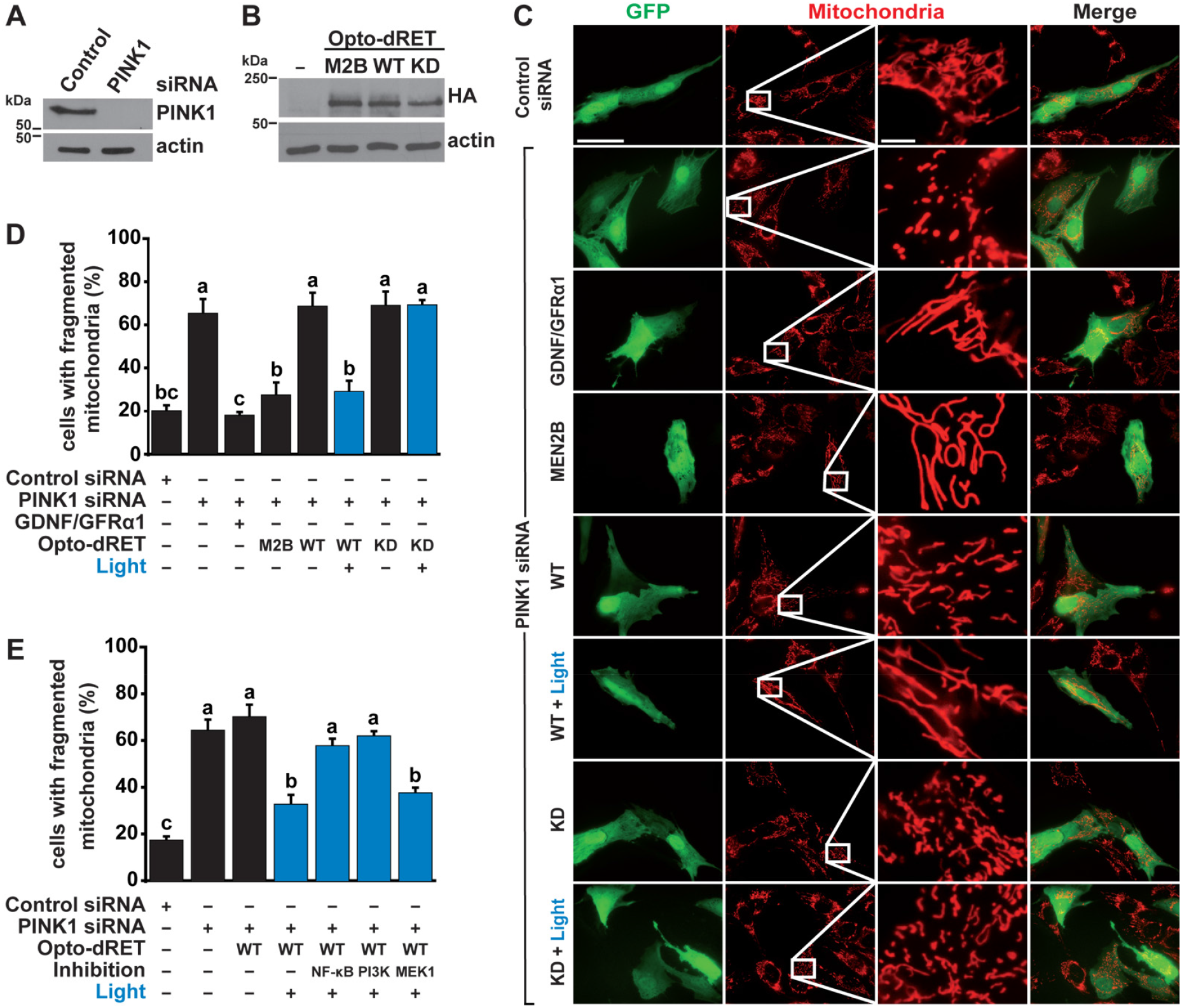
Rescue of mitochondrial fragmentation in human cells. (**A** and **B**) WB analysis of *PINK1* knock-down by siRNA and Opto-dRET expression. (**C**) Representative images for fragmentation of mitochondria induced by *PINK1* silencing. Red: MitoTracker. Green: GFP marker. Scale bar: 200 (column 1, 2) or 20 μm (columns 3, 4). (**D**) Quantification of mitochondrial fragmentation upon stimulation of RET, Opto-dRET, Opto-dRET^MEN2B^ or Opto-dRET^KD^. (**E**) Quantification analysis of mitochondrial fragmentation upon light activation of Opto-dRET and inhibition of NF-кB signaling (by IκB-2S/A), PI3K (by LY294002) or MEK1 (by PD98059). For D and E, mean ± SD for five independent experiments is given (at least 150 cells per condition in each experiment). Means sharing the same label are not significantly different (ANOVA/Bonferroni corrected t-tests, p>.04).

## Discussion

Choreographed signaling of GFs and their cognate RTKs underlies tissue morphogenesis and homeostasis, whereas their aberrant activity is linked to human disorders. For instance, in the case of RET, gain-of-function is implicated in several forms of cancer and loss-of-function is linked to developmental disorders and neurodegeneration (72, 73). Motivated by the importance of RET, we engineered human and *Drosophila* RET receptors that can be activated by light. Recent work has demonstrated light activation of RTKs generally following seminal designs that built on either dimerizing (14) or oligomerizing (15) photoreceptor domains. We developed Opto-hRET and Opto-dRET using the homodimerizing AU1-LOV domain, and whilst LOV domains have enabled dimerization of isolated kinase domains in the past, we here show that this approach is suited for activation of full-length RET receptors. This suggests that enforced association at the RET C-terminus can overcome autoinhibition by elements of the extracellular domain that can counteract ligand-independent dimerization. These light-activated human and *Drosophila* RTKs add to an optogenetic arsenal that already consists of light-activated enzymes and optically-recruited signaling proteins, some of which have already permitted the optogenetic control of cell behavior in *Drosophila* (1, 2).

In the first experiment, we applied Opto-dRET in the *Drosophila* retina to interfere with tissue morphogenesis. Retina development depends on concerted cell proliferation and differentiation events, and RTKs play a key role in ommatidia formation and ommatidial lattice generation. We observed retina malformations specifically in flies that were illuminated during tissue formation stages, in agreement with earlier observations that downstream pathways are not operating at maximal levels during retina development because of multiple and reiterative uses of RTKs (54, 55, 74). This experiment complements recent optogenetic studies in *Drosophila* tissues other than the retina that have incorporated spatio-temporal regulation to identify tissues and stages with high sensitivity to ectopic signals (75–78).

We then explored if optogenetics can suppress phenotypes in a genetic model of PD. Previous optogenetic studies in the context of neurological and neurodegenerative disorders focused on understanding or correcting aberrant electrical activity in excitable cells (7, 8), whilst our goal was the delivery of trophic effects through the optical control of biochemical pro-survival pathways. Our model was *Drosophila* with loss of PINK1, a Ser/Thr kinase that causes autosomal recessive PD (33–35). Although evidently not able to recapitulate all features of human PD, we chose *Drosophila* as the model because PINK1 loss-of-function manifests in robust phenotypes that have previously helped to delineate pathways implicated in mitochondrial physiology and in PD pathogenesis (61, 64, 68, 79–86). Cell degeneration in this model occurs most strongly in cells outside of the nervous system, such as in IFMs, likely because of their high energy demand. Activation of Opto-dRET resulted in efficient suppression of mitochondrial alterations, tissue degeneration and attendant locomotion fitness. We also demonstrated rescue of mitochondrial morphology in PINK1-deficient human cells, and this second model allowed us to identify signaling pathways downstream of dRET that are essential for reversion of the defects. PI3K and NF-кB activity were required to reestablish the healthy mitochondrial network. These pathways are known to act as an important node of crosstalk downstream of tyrosine kinases (87–89), and their involvement is in line with previous observations that the protein networks regulated by GFL/RET overlap with those altered in PD (69, 71). We noted that PINK1 deficiency phenotypes in flies and human cells were only modified by Opto-dRET upon stimulation with light but not in the dark, indicating little background activity of the receptor in the absence of activation of the introduced optical switch. It will be interesting to determine in future studies whether Opto-dRET is efficacious in ameliorating phenotypes in mammalian *in vivo* models of PD.

The new ability to remotely and spatio-temporally control cellular events relevant to human disease has previously inspired the pursuit of optogenetics-based treatment strategies (see above); but what makes optogenetics an attractive method for pro-survival signal delivery, in general or in the context of PD models? The protection or regeneration of cells is a key target in the treatment of a variety of disorders, but the practical application of GFs is challenging. Many GF receptors have widespread tissue distribution and thus systemic growth factor administration may result in off-target effects, such as toxicity or undesired proliferation, in cells other than those targeted (90, 91). Furthermore, many GFs exhibit limited half-life in the circulation or do not reach target tissues (92, 93). Additionally, GF gene therapy results in permanent hyperactivation of signaling pathways that can be linked to side effects and potential safety issues (94). In PD specifically it is not clear if the cellular signaling machinery of degenerating DA neurons can respond to GFLs, e.g. because of impaired RTK retrograde trafficking or expression (23, 95). Optogenetics has properties that may address some of these challenges. For instance, optogenetic control can be reversible and limited to specific cell populations. In addition, optogenetic receptors do not rely on ligand binding in neuronal projections. It has been recently demonstrated that RET downregulation in a mouse model of PD can be compensated by a virally delivered of RET (96). This finding provides an encouraging basis for further exploration of Opto-RETs in mammalian models of PD. Translation of optogenetics into the brain may be facilitated by innovations that are currently pursued by many groups, such as wirelessly-powered microscale electronics that are implantable and biocompatible, or transcranial energy delivery. In this study, we demonstrated in a genetic model of PD that ligand-independent optical delivery of trophic signals is in principle possible, paving the way for future studies in animal models of PD and potentially also other disorders linked to the GF-RTK axis.

## Author contributions (CRediT taxonomy)

Conceptualization, A.I.P., D.S., P.S., K.W. and H.J.; Funding Acquisition, D.S., P.S., K.W. and H.J.; Methodology, A.I.P., P.S., K.W. and H.J.; Project Administration, H.J.; Investigation, A.I.P., N.F., N.H., M.P., E.R. and V.Z.; Data curation, A.I.P., N.F., S.C., N.H., M.P.; Resources, A.G. and J.B.; Supervision, D.S., P.S., K.W. and H.J.; Writing - original Draft, A.I.P. and H.J.; Writing - Review and Editing, D.S., P.S., K.W. and H.J.

## Acknowledgements

We thank R. Cagan, A. Whitworth and J. Nagpal for fly lines and advice, S. Herlitze for provision of a tissue culture illuminator, and Verian Bader for help with statistical analysis. Stocks obtained from the Bloomington *Drosophila* Stock Center (NIH P40OD018537) were used in this study. This work depended on information provided by FlyBase. The research leading to these results has received funding from the People Programme (Marie Curie Actions) of the European Union’s Seventh Framework Programme (FP7/2007-2013) under REA grant agreements 303564 and 334077 (to HJ and DS), from the German Research Foundation (DFG; Research Unit 2848 to KFW; SO1379/4-1 and SO1379/2-1/2-2 to PS), and from the German Academic Exchange Service in The Australia–Germany Joint Research Cooperation Scheme (DAAD, Project ID 57446392, to KFW). A.I.P. was supported by a Ramón Areces post-doctoral fellowship. The Australian Regenerative Medicine Institute is supported by grants from the State Government of Victoria and the Australian Government. The EMBL Australia Partnership Laboratory (EMBL Australia) is supported by the National Collaborative Research Infrastructure Strategy (NCRIS) of the Australian Government.

## Materials and Methods

### Engineering light-activated RET receptors

The gene encoding full-length dRET with the MEN2B M955T substitution (*dRET^MEN2B^*, a kind gift of Ross Cagan, Icahn School of Medicine at Mount Sinai, NY) was amplified from an expression vector by PCR and inserted into pUAST. To obtain *Opto-dRET^MEN2B^* in pUAST, *AU1-LOV* (14) was inserted at the far C-terminus of the receptor. To obtain *Opto-dRET* in pUAST, the M955T substitution of *dRET^MEN2B^* was reverted using site-directed mutagenesis. To express Opto-dRET in mammalian cells, the gene was amplified by PCR and sub-cloned into pcDNA3.1(-) including a hemagglutinin (HA)-epitope. To obtain *Opto-dRET^MEN2B^* in pcDNA3.1(-), the M955T substitution was introduced using site-directed mutagenesis. To obtain the KD variant, the K805M substitution was introduced using site-directed mutagenesis. To express Opto-hRET in mammalian cells, the full-length receptor was inserted into a pcDNA3.1(-) vector containing *AU1-LOV* (14). All constructs were verified by DNA sequencing. Sequences of the receptors are given in **Tables S1** and **S2**.

### Cell culture, transfection and MAPK/ERK pathway activation (HEK293)

The MAPK/ERK pathway was assayed in HEK293 cells with the Elk1 *trans*-reporting system (PathDetect, Agilent). HEK293 cells were maintained in DMEM supplemented with 10% FBS, 100 U/ml penicillin and 0.1 mg/ml streptomycin in a humidified incubator (37°C, 5% CO_2_). 50’000 cells were seeded in each well of white clear bottom 96-well plates (triplicates for each construct) coated with poly-*L*-ornithine (Sigma). Cells were reverse transfected with 3 to 25 ng receptor vector and ~200 ng combined reporter vectors per well using polyethylenimine (Polysciences). Six h after transfection, medium was replaced with CO_2_-independent reduced serum starve medium (Gibco/Life Technologies; supplemented with 0.5% FBS, 2 mM L-Glutamine, 100 U/ml penicillin and 0.1 mg/ml streptomycin). Cells were then either illuminated with blue light in a custom incubator (PT2499, ExoTerra) for 8 h or protected from light with foil as described previously (97). After incubation, plates were processed with a luciferase assay (One-Glo, Promega) and luminescence was detected in a microplate reader (Synergy H1, BioTek). Low-light stimulation (**Figure S3B**) was performed as previously described (47) using a light blocking sample with an optical density of 1 and an external light intensity of 15 μW/cm^2^ (resulting in a final intensity of 1.5 μW/cm^2^).

### Cell culture, RNA interference, transfection and treatments (SH-SY5Y cells)

SH-SY5Y cells (DSMZ ACC 209) were maintained in DEMEM/F-12 (1:1) supplemented with 15% FBS (Sigma), 1% non-essential amino acid solution, 100 U/ml penicillin and 100 μg/ml streptomycin (Life Technologies) in a humidified incubator (37°C, 5% CO_2_). 1.5 × 10^5^ cells were seeded in each well of a 6-well plate containing two 15 mm coverslips per well. Transient co-transfection of siRNA oligos and DNA plasmids were performed using Lipofectamine 2000 (Thermo Fisher). The following three *PINK1* siRNAs were used at a final concentration of 60 pmol/ml each: siRNA *PINK1* HSS127945/127946/185707 (Life Technologies). To identify transfected cells by fluorescence microscopy, a plasmid encoding GFP was co-transfected (0.2 μg/well; in total, 1.2 μg/well were transfected). Mitochondrial morphology was analyzed 2 days after transfection as described below. For illumination of the cells, the 6-well plate was placed in a LED illumination unit inside the incubator. Cells were illuminated for 4 h at a wavelength of 470 nm and an intensity of 232 μW/cm^2^. To activate endogenous RET, cells were treated with recombinant human GDNF (Shenandoah Biotechnology) and human GFRα1 (R&D Systems) for 4 h at a final concentration of 100 ng/ml. Signaling pathway inhibitors were added to cells 1 h prior to illumination at the following concentrations: 50 μM LY294002 (PI3K inhibitor, Cell Signaling) or 30 μM PD98059 (MEK1 inhibitor, Cell Signaling). The HA-IκB-2S32A/S36A plasmid (IκB-2S/A (98)) was generated by overlap extension PCR using the following primers: mut-IκB-2S-fwd CCACGACGCCGGCCTGGACGCCATGAAAG, mut-IκB-2S-rev CGTCTTTCATGGCGTCCAGGCCGGCGTCG, BamHI-IκB2S-fwd ATATGGATCCTTCCAGGCGGCCGAGCGCCCCCAGGAG and IκB2S-NotI-rev ATATGCGGCCGCCTATAACGTCAGACGCTGGCCTCCAAACACACAGTC. The amplified fragments were digested with BamHI and NotI and cloned into the pcDNA3.1-N-HA vector. pEGFP-N3 was purchased from Clontech.

### Analysis of mitochondrial morphology (SH-SY5Y cells)

Mitochondria in SH-SY5Y cells growing on 15 mm coverslips were stained for 15 min with 25 nM MitoTracker red CMXRos (Life Technologies) diluted in cell culture media and then washed twice with medium. Mitochondrial morphology of living cells was immediately analyzed with a fluorescence microscope (Nikon Eclipse E400). Cells displaying an intact network of tubular mitochondria were classified as tubular. When this network was disrupted and mitochondria appeared either globular or rod-like, they were classified as fragmented (70). For quantification of mitochondrial morphology, five independent experiments were performed. At least 150 transfected cells were analyzed per condition for each experiment.

### WB analysis (SH-SY5Y cells)

SH-SY5Y cells were analyzed two days after transient transfection for expression of Opto-dRET constructs and *PINK1* silencing efficiency. For stabilization of endogenous full-length PINK1, cells were treated with 10 μM FCCP (Agilent) for 2 h before cell lysis. Proteins were detected by WB using a monoclonal rabbit anti-PINK1 antibody (1:1000; Cell Signaling, D8G3) or an anti-HA antibody (1:1000; Covance, 16B12) for the Opto-dRET constructs. Data were normalized to monoclonal mouse anti-β-actin staining (1:2000; Sigma, AC-74).

### Fly strains, maintenance and scoring

Flies were raised on standard agar, cornmeal and molasses substrate supplemented with 1.5% nipagin. *GMR-GAL4* flies were a kind gift of Ross Cagan. *PINK1^B9^/FM6; MEF2-GAL4* flies were a gift of Alex Whitworth (University of Cambridge, UK). Transgenic flies expressing Opto-dRET and Opto-dRET^MEN2B^ were generated by injection of pUAST receptor constructs (BestGene). For selection, balanced fly lines (~12 transformants/line) were crossed with *GMR-GAL4* flies. Approximately 10 days after crossing, descendants were visually inspected for the presence of a rough retina phenotype. Rough retina and hollow thorax phenotypes were scored on a dissecting microscope equipped with a digital camera (M205FA and DFC3000G, Leica Microsystems). Genotypes of fly lines utilized in this study are summarized in **Table S3**.

### Light stimulation of flies

Flies were illuminated inside their vials in the custom LED incubator (**Figure S3A**) set to the temperature and light intensities indicated in the text and figures. Vials containing control flies were wrapped with foil and placed in the same incubator. Light incubators were placed in a controlled environment to maintain humidity at 65%. Experiments with GAL4 drivers were conducted at 29°C to increase receptor expression.

### Scanning electron microscopy

Adult flies were anaesthetized with CO_2_, placed in 25% ethanol for 24 h at room temperature (RT) and dehydrated in a graded ethanol series for 3 days. Samples were dried from 100% ethanol with a critical point dryer (EM-CPD300, Leica Microsystems), gold-coated using a sputter coater (EM-ACE600, Leica Microsystems) and imaged at a magnification of 152X (FE-SEM Merlin compact VP, Carl Zeiss; operated at 3 kV).

### Quantification of rough retina phenotype

Three analysis methods were applied to retinas. Fused retinal area was defined as the ratio of the total retina area divided by the total disrupted area. The disrupted area was defined as a region containing two or more fused ommatidia. The number of distinct structures was determined using a distortion algorithm (99). The output of the algorithm is mapping of boundaries surrounding single or fused ommatidia that are the structures of interest. Structure count and structure centers were then identified in Fiji. Regularity was determined based on structure centers and their nearest neighbor distance distributions using macros written in Igor Pro (Wavemetrics). Regularity was defined as the ratio of the mean nearest neighbor distance and its SD for each image (59).

### Transmission electron microscopy

Thoraces were fixed in 2.5% glutaraldehyde and 2% paraformaldehyde in 0.1 M phosphate buffer (pH 7.4) for 2 h at RT. Samples were post-fixed and contrast enhanced with 1% osmium tetroxide in phosphate buffer for 1.5 h and 1% uranyl acetate in 50% ethanol/water for 45 min. Samples were then dehydrated in a graded ethanol series and embedded in Durcupan (Sigma-Aldrich). Ultrathin sections (70 nm) were sliced using a microtome (EM UC7 Ultramicrotome, Leica Microsystems) and mounted on formvar coated copper slot grids. Images were acquired at a magnification of 9000X (Tecnai 10, FEI/Thermo Fisher Scientific; operated at 80 kV, equipped with OSIS Megaview III camera). The electron dense fraction of the cytoplasm (mitochondria) was determined by manual selection, application of threshold and the area fraction command in Fiji.

### ATP determination

Thoraces from two-day old flies were homogenized in 50 μl of extraction buffer (100 mM Tris-HCl, 4 mM EDTA, pH 7.8) supplemented with 6 M Guanidine-HCl using a pellet homogenizer (47747-370, VWR international). The lysate was boiled for 3 min and cleared by centrifugation at 20’000 g for 5 min. The samples were diluted 1:100 in extraction buffer before quantification using a luciferase-based ATP kit (A22066, Thermo Fisher Scientific). Values were expressed relative to total protein concentration measured by using a BCA assay (Pierce). Luminescence and absorbance at 562 nm were measured using the microplate reader. ATP levels were normalized to those of *Opto-dRET* female flies.

### Negative geotaxis (climbing) assay

Male flies of the indicated genotype have been exposed to blue light (I = 320 μW/cm^2^, λ = 470 nm) or kept in the dark during the indicated developmental time points. For each experiment, males hatching on the same day were pooled. Adults were aged for 2-3 days on standard fly food. On day 3, flies were anaesthetized briefly with CO_2_ and 10 flies each were placed in an acrylic glass tube of 30 cm length closed with a flyplug (Carl Roth PK13.1). Flies were allowed to recover and adapt for 1 h. Negative geotaxis climbing performance was then assayed as previously described (100). Flies were tapped down and the number of flies reaching the 15 cm mark within 15 s was recorded. 10 technical repeats (1 min break between repeats) were performed for each genotype to obtain an average climbing score (defined as fly count above the 15 cm line / total fly count). For each genotype and condition, at least 5 independent experiments were performed.

### Statistical analysis

HEK293 cell assays were performed in triplicate wells and in at least three independent experiments. Statistical analysis was performed using unpaired, two-tailed t-tests for comparison of dark and light conditions.

Fly assays were performed in at least three independent experiments with the number of flies indicated in the figures. Statistical analysis of numerical outcomes was performed using one-way analysis of variance (ANOVA) followed by Bonferroni corrected multiple t-test comparisons. For categorical outcomes (thorax defect and climbing pass), SEMs shown in the figure derived from binomial distributions. Statistical significance was tested using Fisher’s exact tests. Climbing experiments were performed in 10 repeats for each animal group consisting of 10 animals. Statistical significance is indicated using the ‘compact letter display’.

SH-SY5Y cell fragmentation assays were performed in five independent experiments with at least 150 cells per condition in each experiment. Statistical analysis was performed using one-way analysis of variance (ANOVA) followed by followed by Bonferroni corrected multiple t-test comparisons. Statistical significance is indicated using ‘compact letter display’.

## Notes

### Competing Interest Statement

The authors have declared no competing interest.

## References

1. H. E. Johnson, J. E. Toettcher, Illuminating developmental biology with cellular optogenetics. Curr Opin Biotechnol 52, 42–48 (2018).

2. G. Guglielmi, H. J. Falk, S. De Renzis, Optogenetic control of protein function: From intracellular processes to tissue morphogenesis. Trends Cell Biol 26, 864–874 (2016).

3. K. Deisseroth, Optogenetics: 10 years of microbial opsins in neuroscience. Nat Neurosci 18, 1213–1225 (2015).

4. L. J. Bugaj et al., Cancer mutations and targeted drugs can disrupt dynamic signal encoding by the Ras-Erk pathway. Science 361(2018).

5. M. Z. Wilson, P. T. Ravindran, W. A. Lim, J. E. Toettcher, Tracing information flow from erk to target gene induction reveals mechanisms of dynamic and combinatorial control. Mol Cell 67, 757–769 e755 (2017).

6. V. Agus, H. Janovjak, Optogenetic methods in drug screening: technologies and applications. Curr Opin Biotechnol 48, 8–14 (2017).

7. J. D. Ordaz, W. Wu, X. M. Xu, Optogenetics and its application in neural degeneration and regeneration. Neural Regen Res 12, 1197–1209 (2017).

8. K. M. Tye, K. Deisseroth, Optogenetic investigation of neural circuits underlying brain disease in animal models. Nat Rev Neurosci 13, 251–266 (2012).

9. V. Gradinaru, M. Mogri, K. R. Thompson, J. M. Henderson, K. Deisseroth, Optical deconstruction of parkinsonian neural circuitry. Science 324, 354–359 (2009).

10. J. A. Steinbeck et al., Optogenetics enables functional analysis of human embryonic stem cell-derived grafts in a Parkinson’s disease model. Nat Biotechnol 33, 204–209 (2015).

11. V. Busskamp et al., Genetic reactivation of cone photoreceptors restores visual responses in retinitis pigmentosa. Science 329, 413–417 (2010).

12. J. B. Bryson et al., Optical control of muscle function by transplantation of stem cell-derived motor neurons in mice. Science 344, 94–97 (2014).

13. T. Bruegmann et al., Optogenetic control of heart muscle in vitro and in vivo. Nat Methods 7, 897–900 (2010).

14. M. Grusch et al., Spatio-temporally precise activation of engineered receptor tyrosine kinases by light. EMBO J 33, 1713–1726 (2014).

15. N. Kim et al., Spatiotemporal control of fibroblast growth factor receptor signals by blue light. Chem Biol 21, 903–912 (2014).

16. V. V. Krishnamurthy et al., Reversible optogenetic control of kinase activity during differentiation and embryonic development. Development 143, 4085–4094 (2016).

17. P. Huang et al., Optical Activation of TrkB Signaling. J Mol Biol 432, 3761–3770 (2020).

18. N. Bunnag et al., An optogenetic method to study signal transduction in intestinal stem cell homeostasis. J Mol Biol 432, 3159–3176 (2020).

19. M. Takahashi, J. Ritz, G. M. Cooper, Activation of a novel human transforming gene, ret, by DNA rearrangement. Cell 42, 581–588 (1985).

20. A. Bjorklund et al., Towards a neuroprotective gene therapy for Parkinson’s disease: use of adenovirus, AAV and lentivirus vectors for gene transfer of GDNF to the nigrostriatal system in the rat Parkinson model. Brain Res 886, 82–98 (2000).

21. C. W. Olanow, R. T. Bartus, L. A. Volpicelli-Daley, J. H. Kordower, Trophic factors for Parkinson’s disease: To live or let die. Movement disorders : official journal of the Movement Disorder Society 30, 1715–1724 (2015).

22. C. Warren Olanow et al., Gene delivery of neurturin to putamen and substantia nigra in Parkinson disease: A double-blind, randomized, controlled trial. Annals of neurology 78, 248–257 (2015).

23. W. J. Marks, Jr. et al., Gene delivery of AAV2-neurturin for Parkinson’s disease: a double-blind, randomised, controlled trial. Lancet Neurol 9, 1164–1172 (2010).

24. F. P. Manfredsson et al., Nigrostriatal rAAV-mediated GDNF overexpression induces robust weight loss in a rat model of age-related obesity. Mol Ther 17, 980–991 (2009).

25. B. Georgievska, D. Kirik, A. Bjorklund, Aberrant sprouting and downregulation of tyrosine hydroxylase in lesioned nigrostriatal dopamine neurons induced by long-lasting overexpression of glial cell line derived neurotrophic factor in the striatum by lentiviral gene transfer. Exp Neurol 177, 461–474 (2002).

26. D. Kirik, C. Rosenblad, A. Bjorklund, R. J. Mandel, Long-term rAAV-mediated gene transfer of GDNF in the rat Parkinson’s model: intrastriatal but not intranigral transduction promotes functional regeneration in the lesioned nigrostriatal system. J Neurosci 20, 4686–4700 (2000).

27. P. Barroso-Chinea et al., Long-term controlled GDNF over-expression reduces dopamine transporter activity without affecting tyrosine hydroxylase expression in the rat mesostriatal system. Neurobiology of disease 88, 44–54 (2016).

28. L. Tenenbaum, M. Humbert-Claude, Glial cell line-derived neurotrophic factor gene delivery in Parkinson’s disease: A delicate balance between neuroprotection, trophic effects, and unwanted compensatory mechanisms. Front Neuroanat 11, 29 (2017).

29. T. M. Axelsen, D. P. D. Woldbye, Gene therapy for Parkinson’s disease, an update. J Parkinsons Dis 8, 195–215 (2018).

30. M. Hahn, J. Bishop, Expression pattern of Drosophila ret suggests a common ancestral origin between the metamorphosis precursors in insect endoderm and the vertebrate enteric neurons. Proc Natl Acad Sci USA 98, 1053–1058 (2001).

31. R. Sugaya, S. Ishimaru, T. Hosoya, K. Saigo, Y. Emori, A Drosophila homolog of human proto-oncogene ret transiently expressed in embryonic neuronal precursor cells including neuroblasts and CNS cells. Mech Dev 45, 139–145 (1994).

32. L. Myers, H. Perera, M. G. Alvarado, T. Kidd, The Drosophila Ret gene functions in the stomatogastric nervous system with the Maverick TGFbeta ligand and the Gfrl co-receptor. Development 145(2018).

33. A. Puschmann et al., Heterozygous PINK1 p.G411S increases risk of Parkinson’s disease via a dominant-negative mechanism. Brain 140, 98–117 (2017).

34. E. M. Valente et al., Hereditary early-onset Parkinson’s disease caused by mutations in PINK1. Science 304, 1158–1160 (2004).

35. E. M. Valente et al., PINK1 mutations are associated with sporadic early-onset parkinsonism. Annals of neurology 56, 336–341 (2004).

36. V. L. Hewitt, A. J. Whitworth, Mechanisms of Parkinson’s disease: lessons from Drosophila. Curr Top Dev Biol 121, 173–200 (2017).

37. A. Voigt, L. A. Berlemann, K. F. Winklhofer, The mitochondrial kinase PINK1: functions beyond mitophagy. J Neurochem 139 Suppl 1, 232–239 (2016).

38. M. Vos, P. Verstreken, C. Klein, Stimulation of electron transport as potential novel therapy in Parkinson’s disease with mitochondrial dysfunction. Biochem Soc Trans 43, 275–279 (2015).

39. S. Schlee, P. Carmillo, A. Whitty, Quantitative analysis of the activation mechanism of the multicomponent growth-factor receptor Ret. Nat Chem Biol 2, 636–644 (2006).

40. S. Jing et al., GDNF-induced activation of the ret protein tyrosine kinase is mediated by GDNFR-alpha, a novel receptor for GDNF. Cell 85, 1113–1124 (1996).

41. N. Asai, T. Iwashita, M. Matsuyama, M. Takahashi, Mechanism of activation of the ret proto-oncogene by multiple endocrine neoplasia 2A mutations. Mol Cell Biol 15, 1613–1619 (1995).

42. B. Freche et al., Inducible dimerization of RET reveals a specific AKT deregulation in oncogenic signaling. J Biol Chem 280, 36584–36591 (2005).

43. R. D. Read et al., A Drosophila model of multiple endocrine neoplasia type 2. Genetics 171, 1057–1081 (2005).

44. C. Abrescia, D. Sjostrand, S. Kjaer, C. F. Ibanez, Drosophila RET contains an active tyrosine kinase and elicits neurotrophic activities in mammalian cells. FEBS Lett 579, 3789–3796 (2005).

45. F. Takahashi et al., AUREOCHROME, a photoreceptor required for photomorphogenesis in stramenopiles. Proc Natl Acad Sci U S A 104, 19625–19630 (2007).

46. T. Toyooka, O. Hisatomi, F. Takahashi, H. Kataoka, M. Terazima, Photoreactions of aureochrome-1. Biophys J 100, 2801–2809 (2011).

47. E. Reichhart, A. Ingles-Prieto, A. M. Tichy, C. McKenzie, H. Janovjak, A phytochrome sensory domain permits receptor activation by red light. Angew Chem Int Ed Engl 55, 6339–6342 (2016).

48. M. J. Kennedy et al., Rapid blue-light-mediated induction of protein interactions in living cells. Nat Methods 7, 973–975 (2010).

49. B. D. Zoltowski, B. Vaccaro, B. R. Crane, Mechanism-based tuning of a LOV domain photoreceptor. Nat Chem Biol 5, 827–834 (2009).

50. K. Sako et al., Optogenetic control of nodal signaling reveals a temporal pattern of nodal signaling regulating cell fate specification during gastrulation. Cell Rep 16, 866–877 (2016).

51. X. Z. Li et al., Identification of a key motif that determines the differential surface levels of RET and TrkB tyrosine kinase receptors and controls depolarization enhanced RET surface insertion. J Biol Chem 287, 1932–1945 (2012).

52. G. Paratcha et al., Released GFRalpha1 potentiates downstream signaling, neuronal survival, and differentiation via a novel mechanism of recruitment of c-Ret to lipid rafts. Neuron 29, 171–184 (2001).

53. M. Takahashi, N. Asai, T. Iwashita, H. Murakami, S. Ito, Mechanisms of development of multiple endocrine neoplasia type 2 and Hirschsprung’s disease by ret mutations. Rec Res Cancer Res 154, 229–236 (1998).

54. N. E. Baker, S. Y. Yu, The EGF receptor defines domains of cell cycle progression and survival to regulate cell number in the developing Drosophila eye. Cell 104, 699–708 (2001).

55. M. Freeman, Reiterative use of the EGF receptor triggers differentiation of all cell types in the Drosophila eye. Cell 87, 651–660 (1996).

56. R. L. Cagan, D. F. Ready, The emergence of order in the Drosophila pupal retina. Dev Biol 136, 346–362 (1989).

57. K. Basler, B. Christen, E. Hafen, Ligand-independent activation of the sevenless receptor tyrosine kinase changes the fate of cells in the developing Drosophila eye. Cell 64, 1069–1081 (1991).

58. J. M. Kramer, B. E. Staveley, GAL4 causes developmental defects and apoptosis when expressed in the developing eye of Drosophila melanogaster. Genet Mol Res 2, 43–47 (2003).

59. H. Wassle, H. J. Riemann, The mosaic of nerve cells in the mammalian retina. Proc R Soc Lond B Biol Sci 200, 441–461 (1978).

60. J. E. Cook, Spatial properties of retinal mosaics: an empirical evaluation of some existing measures. Vis Neurosci 13, 15–30 (1996).

61. J. Park et al., Mitochondrial dysfunction in Drosophila PINK1 mutants is complemented by parkin. Nature 441, 1157–1161 (2006).

62. D. Wang et al., Antioxidants protect PINK1-dependent dopaminergic neurons in Drosophila. Proc Natl Acad Sci U S A 103, 13520–13525 (2006).

63. Y. Yang et al., Mitochondrial pathology and muscle and dopaminergic neuron degeneration caused by inactivation of Drosophila Pink1 is rescued by Parkin. Proc Natl Acad Sci U S A 103, 10793–10798 (2006).

64. I. E. Clark et al., Drosophila pink1 is required for mitochondrial function and interacts genetically with parkin. Nature 441, 1162–1166 (2006).

65. G. Ranganayakulu, D. A. Elliott, R. P. Harvey, E. N. Olson, Divergent roles for NK-2 class homeobox genes in cardiogenesis in flies and mice. Development 125, 3037–3048 (1998).

66. Y. Y. Lin et al., Three-wavelength light control of freely moving Drosophila Melanogaster for less perturbation and efficient social-behavioral studies. Biomed Opt Express 6, 514–523 (2015).

67. M. Hori, K. Shibuya, M. Sato, Y. Saito, Lethal effects of short-wavelength visible light on insects. Sci Rep 4, 7383 (2014).

68. J. C. Greene et al., Mitochondrial pathology and apoptotic muscle degeneration in Drosophila parkin mutants. Proc Natl Acad Sci U S A 100, 4078–4083 (2003).

69. P. Klein et al., Ret rescues mitochondrial morphology and muscle degeneration of Drosophila Pink1 mutants. EMBO J 33, 341–355 (2014).

70. A. K. Lutz et al., Loss of parkin or PINK1 function increases Drp1-dependent mitochondrial fragmentation. J Biol Chem 284, 22938–22951 (2009).

71. D. P. Meka et al., Parkin cooperates with GDNF/RET signaling to prevent dopaminergic neuron degeneration. The Journal of clinical investigation 125, 1873–1885 (2015).

72. L. M. Mulligan, RET revisited: expanding the oncogenic portfolio. Nat Rev Cancer 14, 173–186 (2014).

73. E. R. Kramer, B. Liss, GDNF-Ret signaling in midbrain dopaminergic neurons and its implication for Parkinson disease. FEBS Lett 589, 3760–3772 (2015).

74. D. T. Miller, R. L. Cagan, Local induction of patterning and programmed cell death in the developing Drosophila retina. Development 125, 2327–2335 (1998).

75. H. E. Johnson et al., The spatiotemporal limits of developmental Erk signaling. Dev Cell 40, 185–192 (2017).

76. G. Guglielmi, J. D. Barry, W. Huber, S. De Renzis, An optogenetic method to modulate cell contractility during tissue morphogenesis. Dev Cell 35, 646–660 (2015).

77. X. Wang, L. He, Y. I. Wu, K. M. Hahn, D. J. Montell, Light-mediated activation reveals a key role for Rac in collective guidance of cell movement in vivo. Nat Cell Biol 12, 591–597 (2010).

78. A. L. Patel et al., Optimizing photoswitchable MEK. Proc Natl Acad Sci U S A 116, 25756–25763 (2019).

79. J. C. Greene, A. J. Whitworth, L. A. Andrews, T. J. Parker, L. J. Pallanck, Genetic and genomic studies of Drosophila parkin mutants implicate oxidative stress and innate immune responses in pathogenesis. Hum Mol Genet 14, 799–811 (2005).

80. J. S. Valadas et al., ER lipid defects in neuropeptidergic neurons impair sleep patterns in parkinson’s disease. Neuron 98, 1155–1169 e1156 (2018).

81. J. J. Lee et al., Basal mitophagy is widespread in Drosophila but minimally affected by loss of Pink1 or parkin. J Cell Biol 217, 1613–1622 (2018).

82. M. Vos et al., Cardiolipin promotes electron transport between ubiquinone and complex I to rescue PINK1 deficiency. J Cell Biol 216, 695–708 (2017).

83. J. H. Pogson et al., The complex I subunit NDUFA10 selectively rescues Drosophila pink1 mutants through a mechanism independent of mitophagy. PLoS Genet 10, e1004815 (2014).

84. V. A. Morais et al., PINK1 loss-of-function mutations affect mitochondrial complex I activity via NdufA10 ubiquinone uncoupling. Science 344, 203–207 (2014).

85. E. S. Vincow et al., The PINK1-Parkin pathway promotes both mitophagy and selective respiratory chain turnover in vivo. Proc Natl Acad Sci U S A 110, 6400–6405 (2013).

86. M. Vos et al., Vitamin K2 is a mitochondrial electron carrier that rescues pink1 deficiency. Science 336, 1306–1310 (2012).

87. B. Kloo et al., Critical role of PI3K signaling for NF-kappaB-dependent survival in a subset of activated B-cell-like diffuse large B-cell lymphoma cells. Proc Natl Acad Sci U S A 108, 272–277 (2011).

88. S. Jeay, S. Pianetti, H. M. Kagan, G. E. Sonenshein, Lysyl oxidase inhibits ras-mediated transformation by preventing activation of NF-kappa B. Mol Cell Biol 23, 2251–2263 (2003).

89. C. S. Shi, J. H. Kehrl, PYK2 links G(q)alpha and G(13)alpha signaling to NF-kappa B activation. J Biol Chem 276, 31845–31850 (2001).

90. H. Takayama et al., Diverse tumorigenesis associated with aberrant development in mice overexpressing hepatocyte growth factor/scatter factor. Proc Natl Acad Sci U S A 94, 701–706 (1997).

91. S. E. Kahn, Incretin therapy and islet pathology: a time for caution. Diabetes 62, 2178–2180 (2013).

92. A. Kharitonenkov et al., FGF-21 as a novel metabolic regulator. The Journal of clinical investigation 115, 1627–1635 (2005).

93. T. F. Zioncheck et al., The pharmacokinetics, tissue localization, and metabolic processing of recombinant human hepatocyte growth factor after intravenous administration in rats. Endocrinology 134, 1879–1887 (1994).

94. R. T. Bartus, M. S. Weinberg, R. J. Samulski, Parkinson’s disease gene therapy: success by design meets failure by efficacy. Mol Ther 22, 487–497 (2014).

95. M. Decressac et al., alpha-Synuclein-induced down-regulation of Nurr1 disrupts GDNF signaling in nigral dopamine neurons. Sci Transl Med 4, 163ra156 (2012).

96. N. Volakakis et al., Nurr1 and retinoid X receptor ligands stimulate ret signaling in dopamine neurons and can alleviate alpha-synuclein disrupted gene expression. J Neurosci 35, 14370–14385 (2015).

97. A. Ingles-Prieto et al., Light-assisted small-molecule screening against protein kinases. Nat Chem Biol 11, 952–954 (2015).

98. S. Sun, J. Elwood, W. C. Greene, Both amino- and carboxyl-terminal sequences within I kappa B alpha regulate its inducible degradation. Mol Cell Biol 16, 1058–1065 (1996).

99. Q. Caudron, C. Lyn-Adams, J. A. D. Aston, B. G. Frenguelli, K. G. Moffat, Quantitative assessment of ommatidial distortion in Drosophila melanogaster. Dros Inf Serv 96, 136–144 (2013).

100. Y. O. Ali, W. Escala, K. Ruan, R. G. Zhai, Assaying locomotor, learning, and memory deficits in Drosophila models of neurodegeneration. J Vis Exp (2011).

