## Supplementary Information for "Optogenetic delivery of trophic signals in a genetic model of Parkinson’s disease"

**SUPPLEMENTAL INFORMATION**

Inglés-Prieto *et al.*

**Contents:**

Figures S1 to S4

Tables S1 to S3

Supplemental References

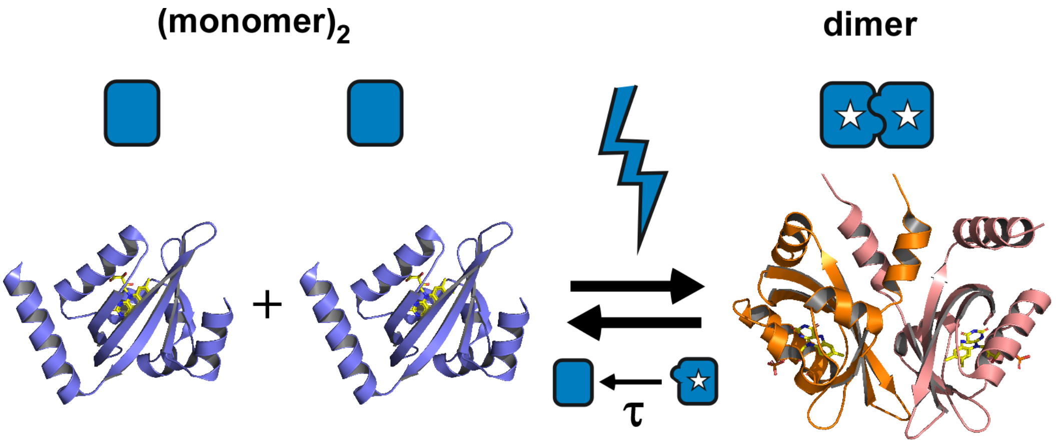


**Figure S1. AU1-LOV is a dimerizing photoreceptor.** Upon photoactivation, AU1-LOV associates in a dimeric 'lit' state (the star denotes the photoadduct state) (Toyooka et al. 2011). Representations of crystal structures obtained for monomeric and dimeric states of AU1-LOV (PDB-IDs: 5DKK and 5DKL; *P. tricornutum*) (Heintz and Schlichting 2016). The lit state AU1-LOV domain relaxes to the dark state with a characteristic lifetime $\tau \sim 600$ s (Mitra et al. 2012; Grusch et al. 2014).

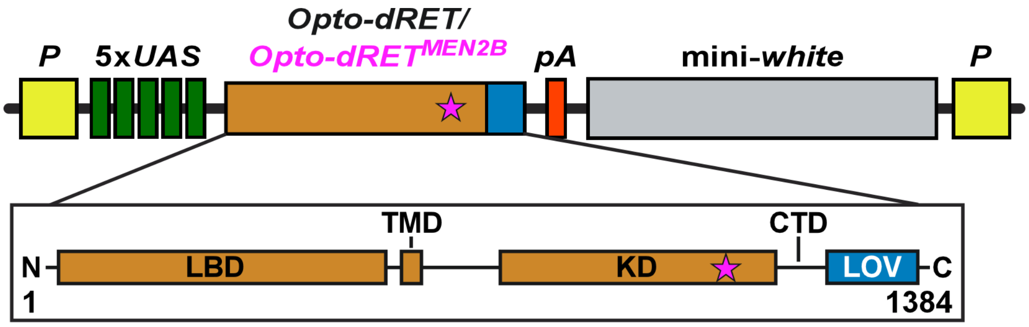


**Figure S2.** ***Opto-dRET*** **construct for GAL4-dependent expression in** ***Drosophila*****.** *Opto-dRET* or *Opto-dRET*$^{MEN2B}$ was inserted in a vector that contains five *UAS* elements, a mini-*white* gene for visualizing transformants and flanking *P*-element terminal repeats. Purple stars represent the kinase domain substitution (M955T; ATG to ACG) in *Opto-dRET*$^{MEN2B}$. N: N-terminus, LBD: extracellular ligand-binding domain, TMD: single-span transmembrane domain, KD: kinase domain, CTD: C-terminal tail domain, LOV: LOV domain, C-terminus.

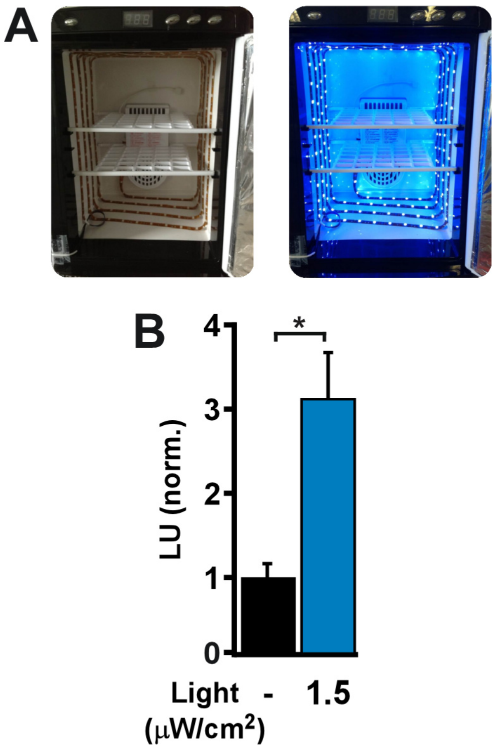


**Figure S3. Low light activation of AU1-LOV.** (**A**) Illumination incubators used in cell and fly experiments. (**B**) Receptor activation in response to blue light stimulation at the indicated intensity for HEK293 cells transfected with *Opto-mFGFR1*. Normalized light units (LU; mean ± SD) for the MAPK/ERK pathway-specific transcriptional reporter in control cells (black) and illuminated cells (blue) are given (n = 9, three independent experiments, t-test, *: p<.0001).

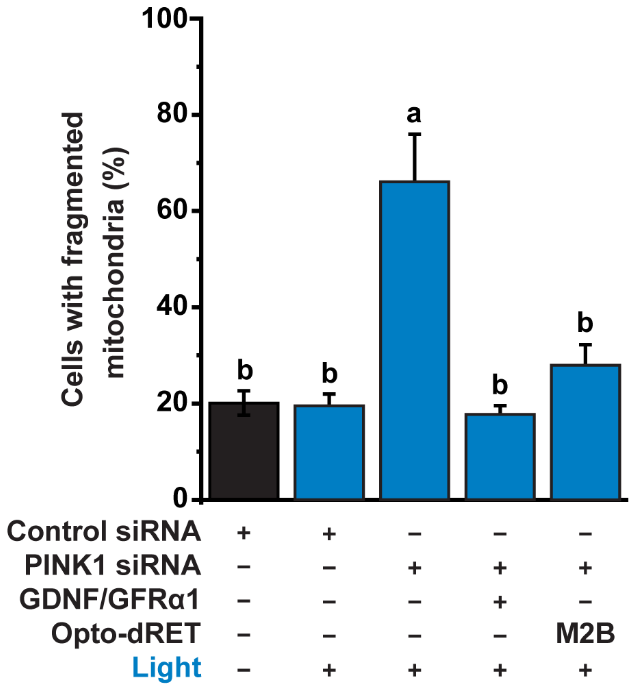


**Figure S4. Light acts specifically through Opto-dRET to rescue mitochondrial fragmentation.** In the absence of Opto-dRET, no effect of blue light (I = 232 μW/cm$^2$, 4h) is observed for cells transfected with control siRNA (compare bars 1 and 2) or *PINK1* siRNA (compare bar 3 of this figure and bar 2 of **Figure 5D**). Likewise, light did not impact the rescue of fragmentation by GDNF/GFRα1 (compare bar 4 of this figure and bar 3 of **Figure 5D**) or by Opto-dRET$^{MEN2B}$ (compare bar 5 of this figure and bar 4 of **Figure 5D**). "M2B" denotes Opto-dRET$^{MEN2B}$. Mean ± SD for five independent experiments is given (>150 cells/condition/experiment). Means sharing the same label are not significantly different (ANOVA/Bonferroni corrected t-tests, p>.04).

**Table S1. (Opto-)dRET sequences.** Uniprot identifiers are given in parentheses.

| Name | Sequence |
|---|---|
| Opto-dRET<br>(*Drosophila* construct) | MESTTIVFVTLLTIITQRKHCAAVDVYFPTTSVKFNLPINEESESIFSKIPL<br>AQFQVLRMEDNRLASDYLYSLEQNPLLRINSSSGEIYMRTDYRSPNSS<br>ATFLVTAFPRDQPDHELLNVSHLSLEVTPQPLEEYCSELEHICFWSSA<br>QYTIAESHGPYRRKDFFEPVLIGALNSRAAKYLCPHVSLEYSLNAGSS<br>HFVLKQNRLYTRQTLDHDELNGLNAKAGQLQARITCTVKLSSRDQRK<br>FSRILDIKLLDRNDNGPKLQESSSKFDFYLEQPYFQADEEAGKKVIYVD<br>KDTLEANAHLVYAVHNDSHGLFRPDCHAYEADHTGRPHTIVSCQLRF<br>SRNGVFRETPYCVSLEARDLTIVSRVDAMSATANVCYHINLSKLHESE<br>QELPQALPLRARQHRIFESEEFNGDSAGRSLSPPTVDYDKDVSVYRS<br>AASNFRVVQPDSFLDLMRLRSIRFDIVEDKLGAFGITSTSGIVFVKNPQ<br>VLEEAPETIYFLNVTWIDQQRLSHVRVINVHLVHGRPENTSCELKVKSR<br>SQTCAQIKYQSQCVRYCGLATGGGSCQWRGSNSAMFGTRYGSCVP<br>ESRYCPDHVCDPLEELNPMACPQDCTPAGRIVGPHSSNENKRGIYSA<br>SGTCICEDNGKCSCAPLDEEPKMKKPRKRKNETEAEPLLGVRRGTPP<br>NQPLQDPMLLGVLNVAGFECDRSCMFFVITCPLLFVLLLLCLLIAQRKM<br>LQRRLGKQSMTTSSKQALPESGGGDFALMPLQSGFRFESGDAKWEF<br>PREKLQLDTVLGEGEFGQVLKGFATEIAGLPGITTVAVKMLKKGSNSV<br>EYMALLSEFQLLQEVSHPNVIKLLGACTSSEAPLLIIEYARYGSLRSYLR<br>LSRKIECAGVDFADGVEPVNVKMVLTFAWQICKGMAYLSELKLVHRDL<br>AARNVLLADGKICKISDFGLTRDVYEDDAYLKRSRDRVPVKWMAPESL<br>ADHVYTSKSDVWSFGVLCWELITLGASPYPGIAPQNLWSLLKTGYRM<br>DRPENCSEAVYSIVRTCWADEPNGRPSFKFLASEFEKLLGNNAKYIDL<br>ETNAVSNPLYCGDDSALITTELGEPESLQHLWSPPKIAYDIHDQATSYD<br>QSEEEMPVTSTAPPGYDLPRPLLDATANGQVLRYENDLRFPLNIRKSS<br>CTPSYSNMTSEPPATTSLPHYSVPVKRGRSYLDMTNKSLIPDNLDSRE<br>FEKHLSKTISFRFSSLLNLSETKEVSPGWQAEDAVTGPDYSLVKALQM<br>AQQNFVITDASLPDNPIVYASRGFLTLTGYSLDQILGRNCRFLQGPETD<br>PRAVDKIRNAITKGVDTSVCLLNYRQDGTTFWNLFFVAGLRDSKGNIV |

| | |
|---|---|
| | NYVGVQSKVSEDYAKLLVNEQNIEYKGVRTSNMLRRKPG |
| dRET (Q7KT06) | MESTTIVFVTLLTIITQRKHCAAVDVYFPTTSVKFNMPINEESESIFSKIP LAQFQVLRMEDNRLASDYLYSLEQNPLLRINSSSGEIYMRTDYRSPNS SATFLVTAFPRDQPDHELLNVSHLSLEVTPQPLEEYCSELEHICFWSS AQYTIAESHGPYRRKDFFEPVLIGALNSRAAKYLCPHVSLEYSLNAGS SHFVLKQNRLYTRQTLDHDELNGLNAKAGQLQARITCTVKLSSRDQR KFSRILDIKLLDRNDNGPKLQESSSKFDFYLEQPYFQADEEAGKKVIYV DKDTLEANAHLVYAVHNDSHGLFRPDCHAYEADHTGRPHTIVSCQLR FSRNGVFRETPYCVSLEARDLTIVSRVDAMSATANVCYHINLSKLHES EQELPQALPLRARQHRIFESEEFNGDSAGRSLSPPTVDYDKDVSVYR SAASNFRVVQPDSFLDLMRLRSIRFDIVEDKLGAFGITSTSGIVFVKNP QVLEEAPETIYFLNVTWIDQQRLSHVRVINVHLVHGRPENTSCELKVK SRSQTCAQIKYQSQCVRYCGLATGGGSCQWRGSNSAMFGTRYGSC VPESRYCPDHVCDPLEELNPMACPQDCTPAGRIVGPHSSNENKRGIY SASGTCICEDNGKCSCAPLDEEPKMKKPRKRKNETEAEPLLGVRRGT PPNQPLQDPMLLGVLNVAGFECDRSCMFFVITCPLLFVLLLLCLLIAQR KMLQRRLGKQSMTTSSKQALPESGGGDFALMPLQSGFRFESGDAKW EFPREKLQLDTVLGEGEFGQVLKGFATEIAGLPGITTVAVKMLKKGSN SVEYMALLSEFQLLQEVSHPNVIKLLGACTSSEAPLLIIEYARYGSLRSY LRLSRKIECAGVDFADGVEPVNVKMVLTFAWQICKGMAYLSELKLVHR DLAARNVLLADGKICKISDFGLTRDVYEDDAYLKRSRDRVPVKWMAPE SLADHVYTSKSDVWSFGVLCWELITLGASPYPGIAPQNLWSLLKTGYR MDRPENCSEAVYSIVRTCWADEPNGRPSFKFLASEFEKLLGNNAKYID LETNAVSNPLYCGDDSALITTELGEPESLQHLWSPPKIAYDIHDQATSY DQSEEEMPVTSTAPPGYDLPRPLLDATANGQVLRYENDLRFPLNIRKS SCTPSYSNMTSEPPATTSLPHYSVPVKRGRSYLDMTNKSLIPDNLDSR EFEKHLSKTISFRFSSLLNLSETKEVSPGWQAEDAV |
| LOV domain of *V. frigida* AUREOCHROME1 (residues 204 to 348 of | PDYSLVKALQMAQQNFVITDASLPDNPIVYASRGFLTLTGYSLDQILGR NCRFLQGPETDPRAVDKIRNAITKGVDTSVCLLNYRQDGTTFWNLFFV AGLRDSKGNIVNYVGVQSKVSEDYAKLLVNEQNIEYKGVRTSNMLRR |

| | |
|---|---|
| A8QW55) (AU1-LOV) | K |
| *Opto-dRET* (cDNA)<br><br>(*Drosophila* construct) | ATGGAGTCAACTACTATTGTTTTTGTGACTCTGCTCACAATTATAAC<br>CCAACGTAAACACTGTGCGGCCGTCGATGTTTACTTTCCCACCACG<br>TCGGTGAAATTCAATCTGCCCATCAATGAGGAATCGGAGAGCATAT<br>TCTCCAAAATCCCGCTAGCCCAGTTCCAAGTGCTGCGGATGGAGG<br>ACAATCGGTTAGCCAGTGATTACTTGTATAGCCTGGAGCAGAATCC<br>ACTACTCCGAATAAACAGTTCCTCCGGCGAGATATATATGCGCACT<br>GACTACCGCTCACCAAACTCAAGTGCCACATTCTTGGTGACCGCAT<br>TTCCCAGAGATCAACCGGATCACGAGCTGCTGAATGTTTCGCATCT<br>TTCGTTGGAGGTTACACCTCAGCCCCTGGAGGAGTACTGTTCGGA<br>ACTGGAGCACATTTGCTTCTGGAGCAGTGCTCAGTACACTATAGCA<br>GAGTCGCACGGTCCATATCGGCGGAAGGATTTTTTTGAACCCGTAC<br>TTATCGGCGCCCTTAATTCCCGCGCTGCGAAGTATCTGTGTCCTCA<br>TGTATCCCTGGAATATTCCCTGAACGCTGGTAGTTCCCATTTTGTTT<br>TGAAACAAAATCGACTCTACACCCGACAAACCTTGGATCACGACGA<br>GCTCAATGGACTGAATGCCAAGGCAGGGCAGCTGCAGGCCAGGAT<br>TACCTGCACGGTTAAATTGTCCAGCAGGGATCAGAGAAAATTCTCG<br>CGCATCTTGGATATCAAGTTACTGGATCGCAATGATAATGGACCCA<br>AGTTGCAGGAGAGTAGCTCTAAGTTTGATTTCTATCTGGAGCAGCC<br>CTACTTCCAAGCGGACGAGGAGGCGGGAAAAAAAGTAATCTACGT<br>GGACAAGGATACATTGGAGGCAAATGCTCACCTTGTCTACGCCGT<br>CCACAATGACTCTCATGGTCTGTTTCGGCCCGACTGCCACGCCTAC<br>GAGGCGGATCACACGGGCAGACCACATACCATCGTCAGTTGTCAA<br>CTGCGATTCTCCCGAAACGGTGTCTTCCGGGAAACCCCCTATTGTG<br>TGTCCTTGGAGGCTCGGGATCTGACCATTGTAAGCCGTGTCGATG<br>CCATGTCAGCGACAGCCAATGTTTGCTATCATATTAATCTGAGTAA<br>GCTTCACGAATCTGAGCAAGAATTACCGCAAGCTCTTCCCCTACGG<br>GCACGTCAACATCGAATATTCGAGAGCGAAGAATTCAATGGAGATT<br>CTGCAGGCCGATCTCTAAGTCCTCCGACCGTGGATTACGATAAGG<br>ATGTTTCCGTATACAGATCGGCTGCTTCTAATTTTCGAGTTGTCCAG |

| | |
|---|---|
| | CCTGACAGTTTTTTGGACTTGATGCGGTTACGATCTATTCGATTCGA<br>TATTGTGGAGGATAAACTTGGAGCTTTTGGTATTACCTCAACATCG<br>GGTATTGTCTTTGTGAAGAACCCACAGGTTTTGGAGGAGGCACCG<br>GAAACCATATACTTCCTGAATGTCACCTGGATCGATCAGCAAAGGC<br>TGTCGCACGTGAGAGTGATCAATGTGCACTTGGTTCATGGTAGACC<br>CGAGAATACTAGTTGCGAACTGAAGGTCAAGTCTCGATCACAGACA<br>TGTGCCCAGATTAAATACCAATCGCAATGCGTTCGATATTGCGGCT<br>TGGCCACAGGTGGTGGATCTTGCCAGTGGAGGGGGTCCAACTCAG<br>CCATGTTCGGCACTAGATATGGTTCCTGTGTACCCGAATCTCGTTA<br>CTGTCCAGATCATGTCTGTGATCCCCTAGAGGAACTGAATCCTATG<br>GCCTGTCCGCAGGATTGCACGCCAGCTGGAAGAATCGTGGGTCCC<br>CATTCAAGTAATGAGAATAAGAGAGGGATATACAGTGCCTCGGGTA<br>CCTGCATTTGCGAGGATAATGGCAAGTGCTCGTGCGCTCCGTTAG<br>ATGAGGAACCCAAGATGAAGAAACCGCGAAAACGAAAAAACGAAA<br>CAGAGGCGGAACCTCTGCTGGGGGTACGAAGGGGCACTCCTCCG<br>AATCAGCCACTTCAGGATCCCATGCTTCTGGGTGTCCTAAATGTGG<br>CCGGTTTCGAATGCGATCGCTCCTGCATGTTCTTCGTGATCACGTG<br>CCCTCTATTGTTCGTTCTCCTGCTCCTCTGTTTGCTGATTGCGCAG<br>AGAAAGATGCTCCAACGTCGCTTGGGCAAGCAATCAATGACCACG<br>TCGTCGAAACAAGCTCTTCCGGAATCAGGAGGCGGAGATTTCGCC<br>TTGATGCCGCTGCAGAGTGGCTTCAGGTTCGAAAGTGGGGATGCC<br>AAATGGGAGTTTCCCAGGGAAAAACTGCAACTAGATACGGTCTTGG<br>GAGAGGGTGAATTTGGTCAGGTGCTAAAGGGCTTTGCCACCGAGA<br>TCGCTGGCTTGCCGGGAATAACCACGGTGGCCGTAAAGATGCTCA<br>AGAAGGGCTCCAATTCAGTGGAGTACATGGCCCTGCTTTCGGAGT<br>TTCAGCTCCTCCAGGAGGTCTCTCACCCGAATGTGATCAAGCTGCT<br>AGGCGCCTGCACCTCCTCCGAAGCACCTCTCCTGATCATCGAGTA<br>TGCTCGGTATGGCTCTCTGAGGAGCTATCTTCGACTCAGTCGGAA<br>GATCGAGTGTGCCGGCGTAGATTTCGCAGATGGAGTGGAGCCTGT<br>TAATGTTAAGATGGTACTTACCTTTGCTTGGCAGATTTGCAAGGGTA |

| | |
|---|---|
| | TGGCTTACCTTTCTGAGTTAAAGTTGGTTCATCGTGATTTGGCTGCT AGAAATGTGCTCCTTGCGGATGGCAAGATATGCAAAATATCAGATT TCGGACTGACTCGAGATGTTTACGAGGACGATGCCTATTTAAAGAG ATCCCGAGATCGTGTGCCCGTCAAGTGGATGGCTCCGGAATCTTT AGCGGATCATGTGTATACCAGCAAATCGGATGTGTGGTCCTTTGGC GTTCTCTGCTGGGAACTAATCACTCTGGGAGCCTCTCCGTATCCTG GCATTGCTCCCCAGAATTTGTGGTCCTTGCTGAAGACGGGCTACC GCATGGACAGACCAGAAAACTGTTCGGAGGCTGTCTACTCTATAGT TCGAACTTGCTGGGCAGACGAGCCAAATGGAAGACCCTCATTCAA GTTTTTAGCATCGGAGTTTGAGAAGCTATTGGGAAACAATGCCAAG TACATAGATCTGGAAACGAATGCCGTTTCGAATCCCCTTTATTGTG GGGATGATTCCGCCTTAATAACCACGGAATTGGGCGAACCAGAAT CGTTGCAGCACCTTTGGTCACCTCCCAAAATAGCCTACGACATCCA TGACCAGGCCACCAGCTACGATCAGTCTGAGGAGGAGATGCCAGT GACTTCAACGGCTCCGCCGGGTTACGACTTACCACGACCTTTGCTT GATGCTACCGCCAACGGGCAGGTTTTGCGATACGAAAACGATTTG CGATTCCCCTTAAATATTCGGAAATCCAGTTGTACTCCAAGCTACA GCAACATGACCAGTGAACCTCCAGCGACCACCTCACTGCCACATTA TTCTGTCCCCGTGAAGAGGGGTCGATCCTACCTGGATATGACCAA CAAGAGTCTCATCCCAGACAACCTGGACAGCAGGGAGTTTGAAAA GCATCTGTCCAAGACCATCTCGTTCCGTTTCTCTAGTTTGCTGAATC TCAGTGAAACGAAGGAGGTGAGTCCAGGATGGCAAGCTGAGGATG CAGTCACCGGTGGACCTGACTACAGTCTCGTGAAGGCTCTGCAAA TGGCACAACAGAATTTTGTCATTACAGACGCCTCCCTCCCAGACAA CCCTATCGTCTACGCCAGTAGAGGGTTTCTGACACTGACAGGCTAT TCTCTCGACCAGATCCTGGGCAGGAACTGCAGGTTTCTGCAAGGG CCAGAAACAGACCCAAGAGCTGTGGATAAGATCAGGAATGCCATC ACCAAAGGCGTTGATACCAGTGTCTGTCTGCTGAATTATAGACAGG ATGGCACAACCTTCTGGAATCTCTTCTTCGTGGCTGGACTCAGAGA TTCTAAGGGCAATATTGTCAACTACGTCGGAGTGCAGTCAAAGGTG |

| | |
|---|---|
| | AGCGAAGATTATGCCAAGCTGCTGGTCAACGAGCAGAACATTGAG<br>TACAAAGGTGTGCGCACCAGTAACATGCTGCGCAGAAAGCCCGGT<br>TAG |

**Table S2. (Opto-)hRET sequences.** Uniprot identifiers are given in parentheses.

| **Name** | **Sequence** |
|---|---|
| Opto-hRET | MAKATSGAAGLRLLLLLLLPLLGKVALGLYFSRDAYWEKLYVDQAAGT<br>PLLYVHALRDAPEEVPSFRLGQHLYGTYRTRLHENNWICIQEDTGLLY<br>LNRSLDHSSWEKLSVRNRGFPLLTVYLKVFLSPTSLREGECQWPGCA<br>RVYFSFFNTSFPACSSLKPRELCFPETRPSFRIRENRPPGTFHQFRLL<br>PVQFLCPNISVAYRLLEGEGLPFRCAPDSLEVSTRWALDREQREKYEL<br>VAVCTVHAGAREEVVMVPFPVTVYDEDDSAPTFPAGVDTASAVVEFK<br>RKEDTVVATLRVFDADVVPASGELVRRYTSTLLPGDTWAQQTFRVEH<br>WPNETSVQANGSFVRATVHDYRLVLNRNLSISENRTMQLAVLVNDSD<br>FQGPGAGVLLLHFNVSVLPVSLHLPSTYSLSVSRRARRFAQIGKVCVE<br>NCQAFSGINVQYKLHSSGANCSTLGVVTSAEDTSGILFVNDTKALRRP<br>KCAELHYMVVATDQQTSRQAQAQLLVTVEGSYVAEEAGCPLSCAVSK<br>RRLECEECGGLGSPTGRCEWRQGDGKGITRNFSTCSPSTKTCPDGH<br>CDVVETQDINICPQDCLRGSIVGGHEPGEPRGIKAGYGTCNCFPEEEK<br>CFCEPEDIQDPLCDELCRTVIAAAVLFSFIVSVLLSAFCIHCYHKFAHKP<br>PISSAEMTFRRPAQAFPVSYSSSGARRPSLDSMENQVSVDAFKILEDP<br>KWEFPRKNLVLGKTLGEGEFGKVVKATAFHLKGRAGYTTVAVKMLKE<br>NASPSELRDLLSEFNVLKQVNHPHVIKLYGACSQDGPLLLIVEYAKYGS<br>LRGFLRESRKVGPGYLGSGGSRNSSSLDHPDERALTMGDLISFAWQI<br>SQGMQYLAEMKLVHRDLAARNILVAEGRKMKISDFGLSRDVYEEDSY<br>VKRSQGRIPVKWMAIESLFDHIYTTQSDVWSFGVLLWEIVTLGGNPYP<br>GIPPERLFNLLKTGHRMERPDNCSEEMYRLMLQCWKQEPDKRPVFA<br>DISKDLEKMMVKRRDYLDLAASTPSDSLIYDDGLSEEETPLVDCNNAP<br>LPRALPSTWIENKLYGRISHAFTRFTGGPDYSLVKALQMAQQNFVITD<br>ASLPDNPIVYASRGFLTLTGYSLDQILGRNCRFLQGPETDPRAVDKIRN<br>AITKGVDTSVCLLNYRQDGTTFWNLFFVAGLRDSKGNIVNYVGVQSKV<br>SEDYAKLLVNEQNIEYKGVRTSNMLRRKPGGSGVDYPYDVPDYALD |
| hRET (P07949-2) | MAKATSGAAGLRLLLLLLLPLLGKVALGLYFSRDAYWEKLYVDQAAGT<br>PLLYVHALRDAPEEVPSFRLGQHLYGTYRTRLHENNWICIQEDTGLLY |

| | |
|---|---|
| | LNRSLDHSSWEKLSVRNRGFPLLTVYLKVFLSPTSLREGECQWPGCA RVYFSFFNTSFPACSSLKPRELCFPETRPSFRIRENRPPGTFHQFRLL PVQFLCPNISVAYRLLEGEGLPFRCAPDSLEVSTRWALDREQREKYEL VAVCTVHAGAREEVVMVPFPVTVYDEDDSAPTFPAGVDTASAVVEFK RKEDTVVATLRVFDADVVPASGELVRRYTSTLLPGDTWAQQTFRVEH WPNETSVQANGSFVRATVHDYRLVLNRNLSISENRTMQLAVLVNDSD FQGPGAGVLLLHFNVSVLPVSLHLPSTYSLSVSRRARRFAQIGKVCVE NCQAFSGINVQYKLHSSGANCSTLGVVTSAEDTSGILFVNDTKALRRP KCAELHYMVVATDQQTSRQAQAQLLVTVEGSYVAEEAGCPLSCAVSK RRLECEECGGLGSPTGRCEWRQGDGKGITRNFSTCSPSTKTCPDGH CDVVETQDINICPQDCLRGSIVGGHEPGEPRGIKAGYGTCNCFPEEEK CFCEPEDIQDPLCDELCRTVIAAAVLFSFIVSVLLSAFCIHCYHKFAHKP PISSAEMTFRRPAQAFPVSYSSSGARRPSLDSMENQVSVDAFKILEDP KWEFPRKNLVLGKTLGEGEFGKVVKATAFHLKGRAGYTTVAVKMLKE NASPSELRDLLSEFNVLKQVNHPHVIKLYGACSQDGPLLLIVEYAKYGS LRGFLRESRKVGPGYLGSGGSRNSSSLDHPDERALTMGDLISFAWQI SQGMQYLAEMKLVHRDLAARNILVAEGRKMKISDFGLSRDVYEEDSY VKRSQGRIPVKWMAIESLFDHIYTTQSDVWSFGVLLWEIVTLGGNPYP GIPPERLFNLLKTGHRMERPDNCSEEMYRLMLQCWKQEPDKRPVFA DISKDLEKMMVKRRDYLDLAASTPSDSLIYDDGLSEEETPLVDCNNAP LPRALPSTWIENKLYGRISHAFTRF |
| *Opto-hRET* (cDNA) | ATGGCGAAGGCGACGTCCGGTGCCGCGGGGCTGCGTCTGCTGTT GCTGCTGCTGCTGCCGCTGCTAGGCAAAGTGGCATTGGGCCTCTA CTTCTCGAGGGATGCTTACTGGGAGAAGCTGTATGTGGACCAGGC GGCCGGCACGCCCTTGCTGTACGTCCATGCCCTGCGGGACGCCC CTGAGGAGGTGCCCAGCTTCCGCCTGGGCCAGCATCTCTACGGCA CGTACCGCACACGGCTGCATGAGAACAACTGGATCTGCATCCAGG AGGACACCGGCCTCCTCTACCTTAACCGGAGCCTGGACCATAGCT CCTGGGAGAAGCTCAGTGTCCGCAACCGCGGCTTTCCCCTGCTCA CCGTCTACCTCAAGGTCTTCCTGTCACCCACATCCCTTCGTGAGGG |

| | |
|---|---|
| | CGAGTGCCAGTGGCCAGGCTGTGCCCGCGTATACTTCTCCTTCTT<br>CAACACCTCCTTTCCAGCCTGCAGCTCCCTCAAGCCCCGGGAGCT<br>CTGCTTCCCAGAGACAAGGCCCTCCTTCCGCATTCGGGAGAACCG<br>ACCCCCAGGCACCTTCCACCAGTTCCGCCTGCTGCCTGTGCAGTT<br>CTTGTGCCCCAACATCAGCGTGGCCTACAGGCTCCTGGAGGGTGA<br>GGGTCTGCCCTTCCGCTGCGCCCCGGACAGCCTGGAGGTGAGCA<br>CGCGCTGGGCCCTGGACCGCGAGCAGCGGGAGAAGTACGAGCTG<br>GTGGCCGTGTGCACCGTGCACGCCGGCGCGCGCGAGGAGGTGGT<br>GATGGTGCCCTTCCCGGTGACCGTGTACGACGAGGACGACTCGG<br>CGCCCACCTTCCCCGCGGGCGTCGACACCGCCAGCGCCGTGGTG<br>GAGTTCAAGCGGAAGGAGGACACCGTGGTGGCCACGCTGCGTGT<br>CTTCGATGCAGACGTGGTACCTGCATCAGGGGAGCTGGTGAGGCG<br>GTACACAAGCACGCTGCTCCCCGGGGACACCTGGGCCCAGCAGA<br>CCTTCCGGGTGGAACACTGGCCCAACGAGACCTCGGTCCAGGCCA<br>ACGGCAGCTTCGTGCGGGCGACCGTACATGACTATAGGCTGGTTC<br>TCAACCGGAACCTCTCCATCTCGGAGAACCGCACCATGCAGCTGG<br>CGGTGCTGGTCAATGACTCAGACTTCCAGGGCCCAGGAGCGGGC<br>GTCCTCTTGCTCCACTTCAACGTGTCGGTGCTGCCGGTCAGCCTG<br>CACCTGCCCAGTACCTACTCCCTCTCCGTGAGCAGGAGGGCTCGC<br>CGATTTGCCCAGATCGGGAAAGTCTGTGTGGAAAACTGCCAGGCA<br>TTCAGTGGCATCAACGTCCAGTACAAGCTGCATTCCTCTGGTGCCA<br>ACTGCAGCACGCTAGGGGTGGTCACCTCAGCCGAGGACACCTCG<br>GGGATCCTGTTTGTGAATGACACCAAGGCCCTGCGGCGGCCCAAG<br>TGTGCCGAACTTCACTACATGGTGGTGGCCACCGACCAGCAGACC<br>TCTAGGCAGGCCCAGGCCCAGCTGCTTGTAACAGTGGAGGGGTCA<br>TATGTGGCCGAGGAGGCGGGCTGCCCCCTGTCCTGTGCAGTCAG<br>CAAGAGACGGCTGGAGTGTGAGGAGTGTGGCGGCCTGGGCTCCC<br>CAACAGGCAGGTGTGAGTGGAGGCAAGGAGATGGCAAAGGGATC<br>ACCAGGAACTTCTCCACCTGCTCTCCCAGCACCAAGACCTGCCCC<br>GACGGCCACTGCGATGTTGTGGAGACCCAAGACATCAACATTTGC |

| | |
|---|---|
| | CCTCAGGACTGCCTCCGGGGCAGCATTGTTGGGGGACACGAGCC<br>TGGGGAGCCCCGGGGGATTAAAGCTGGCTATGGCACCTGCAACTG<br>CTTCCCTGAGGAGGAGAAGTGCTTCTGCGAGCCCGAAGACATCCA<br>GGATCCACTGTGCGACGAGCTGTGCCGCACGGTGATCGCAGCCG<br>CTGTCCTCTTCTCCTTCATCGTCTCGGTGCTGCTGTCTGCCTTCTG<br>CATCCACTGCTACCACAAGTTTGCCCACAAGCCACCCATCTCCTCA<br>GCTGAGATGACCTTCCGGAGGCCCGCCCAGGCCTTCCCGGTCAG<br>CTACTCCTCTTCCGGTGCCCGCCGGCCCTCGCTGGACTCCATGGA<br>GAACCAGGTCTCCGTGGATGCCTTCAAGATCCTGGAGGATCCAAA<br>GTGGGAATTCCCTCGGAAGAACTTGGTTCTTGGAAAAACTCTAGGA<br>GAAGGCGAATTTGGAAAAGTGGTCAAGGCAACGGCCTTCCATCTG<br>AAAGGCAGAGCAGGGTACACCACGGTGGCCGTGAAGATGCTGAAA<br>GAGAACGCCTCCCCGAGTGAGCTTCGAGACCTGCTGTCAGAGTTC<br>AACGTCCTGAAGCAGGTCAACCACCCACATGTCATCAAATTGTATG<br>GGGCCTGCAGCCAGGATGGCCCGCTCCTCCTCATCGTGGAGTAC<br>GCCAAATACGGCTCCCTGCGGGGCTTCCTCCGCGAGAGCCGCAAA<br>GTGGGGCCTGGCTACCTGGGCAGTGGAGGCAGCCGCAACTCCAG<br>CTCCCTGGACCACCCGGATGAGCGGGCCCTCACCATGGGCGACC<br>TCATCTCATTTGCCTGGCAGATCTCACAGGGGATGCAGTATCTGGC<br>CGAGATGAAGCTCGTTCATCGGGACTTGGCAGCCAGAAACATCCT<br>GGTAGCTGAGGGGCGGAAGATGAAGATTTCGGATTTCGGCTTGTC<br>CCGAGATGTTTATGAAGAGGATTCCTACGTGAAGAGGAGCCAGGG<br>TCGGATTCCAGTTAAATGGATGGCAATTGAATCCCTTTTTGATCATA<br>TCTACACCACGCAAAGTGATGTATGGTCTTTTGGTGTCCTGCTGTG<br>GGAGATCGTGACCCTAGGGGGAAACCCCTATCCTGGGATTCCTCC<br>TGAGCGGCTCTTCAACCTTCTGAAGACCGGCCACCGGATGGAGAG<br>GCCAGACAACTGCAGCGAGGAGATGTACCGCCTGATGCTGCAATG<br>CTGGAAGCAGGAGCCGGACAAAAGGCCGGTGTTTGCGGACATCA<br>GCAAAGACCTGGAGAAGATGATGGTTAAGAGGAGAGACTACTTGG<br>ACCTTGCGGCGTCCACTCCATCTGACTCCCTGATTTATGACGACGG |

| | |
|---|---|
| | CCTCTCAGAGGAGGAGACACCGCTGGTGGACTGTAATAATGCCCC<br>CCTCCCTCGAGCCCTCCCTTCCACATGGATTGAAAACAAACTCTAT<br>GGTAGAATTTCCCATGCATTTACTAGATTCAccggTGGACCTGACTA<br>CAGTCTCGTGAAGGCTCTGCAAATGGCACAACAGAATTTTGTCATT<br>ACAGACGCCTCCCTCCCAGACAACCCTATCGTCTACGCCAGTAGA<br>GGGTTTCTGACACTGACAGGCTATTCTCTCGACCAGATCCTGGGCA<br>GGAACTGCAGGTTTCTGCAAGGGCCAGAAACAGACCCAAGAGCTG<br>TGGATAAGATCAGGAATGCCATCACCAAAGGCGTTGATACCAGTGT<br>CTGTCTGCTGAATTATAGACAGGATGGCACAACCTTCTGGAATCTC<br>TTCTTCGTGGCTGGACTCAGAGATTCTAAGGGCAATATTGTCAACT<br>ACGTCGGAGTGCAGTCAAAGGTGAGCGAAGATTATGCCAAGCTGC<br>TGGTCAACGAGCAGAACATTGAGTACAAAGGTGTGCGCACCAGTA<br>ACATGCTGCGCAGAAAGCCCGGTGGATCCGGAGTCGACTATCCGT<br>ACGACGTACCAGACTACGCACTCGACTAA |

**Table S3. Genotypes of flies in this study.**

| | SEX CHR | CHR 2 | CHR 3 |
|---|---|---|---|
| Figure 2B,H-I | | *UAS-dRET::AU1LOV/+* | |
| Figure 2C,H-I | | *GMR-GAL4/+* | *UAS_dRET*$^{MEN2B}$*::AU1LOV/+* |
| Figure 2D,E,H-I | | *GMR-GAL4/+* | |
| Figure 2F-I | | *GMR-GAL4/UAS-dRET::AU1LOV* | |
| Figure 3B | *PINK1*$^{B9}$*/+* | | *MEF2-GAL4/+* |
| Figure 3B,C | *PINK1*$^{B9}$*/Y* | | *MEF2-GAL4/+* |
| Figure 3B,C | *PINK1*$^{B9}$*/Y* | *UAS-dRET::AU1LOV/+* | *MEF2-GAL4/+* |
| Figure 3B | *PINK1*$^{B9}$*/Y* | | *MEF2-GAL4/UAS_dRET*$^{MEN2B}$*::AU1LOV* |
| Figure 3C | *+/Y* | | |
| Figure 4A,B,F | *PINK1*$^{B9}$*/+* | | *MEF2-GAL4/+* |
| Figure 4A,C,F | *PINK1*$^{B9}$*/Y* | | *MEF2-GAL4/+* |
| Figure 4A,D-F | *PINK1*$^{B9}$*/Y* | *UAS-dRET::AU1LOV/+* | *MEF2-GAL4/+* |

**Supplemental References**


Grusch, M., Schelch, K., Riedler, R., Reichhart, E., Differ, C., Berger, W., Ingles-Prieto, A., Janovjak, H., 2014. Spatio-temporally precise activation of engineered receptor tyrosine kinases by light. EMBO J 33, 1713-1726.

Heintz, U., Schlichting, I., 2016. Blue light-induced LOV domain dimerization enhances the affinity of Aureochrome 1a for its target DNA sequence. Elife 5, e11860.

Mitra, D., Yang, X., Moffat, K., 2012. Crystal structures of Aureochrome1 LOV suggest new design strategies for optogenetics. Structure 20, 698-706.

Toyooka, T., Hisatomi, O., Takahashi, F., Kataoka, H., Terazima, M., 2011. Photoreactions of aureochrome-1. Biophys J 100, 2801-2809.
